# Complete sequencing of the mouse pseudoautosomal region, the most rapidly evolving ‘chromosome’

**DOI:** 10.1101/2022.03.26.485930

**Authors:** Takaoki Kasahara, Kazuyuki Mekada, Kuniya Abe, Alan Ashworth, Tadafumi Kato

## Abstract

The pseudoautosomal region (PAR) of mammalian sex chromosomes is a small region of sequence identity that allows pairing, crossover, recombination, and proper chromosome segregation during male meiosis. The structure of the mouse PAR is largely unknown. Here, we developed a new assembly method to robustly resolve repetitive sequences and employed highly accurate long-read sequencing data to reveal the entire PAR sequence. The PAR of the widely-used inbred strain C57BL/6J is ∼700 kb, comprising 10 protein-coding genes in a mass of complex repetitive sequences. A large segmental duplication exhibiting copy-number polymorphisms even among C57BL/6J littermates is present. High GC-content exons and short introns are common properties of PAR genes and are the consequence of maintaining gene function, while PAR is rapidly evolving. Elucidating the mouse PAR sequence completes the mouse euchromatic genome sequencing and enables the exploration of the function and evolution of the PAR using modern molecular genetic approaches.

## Introduction

The sex chromosomes (X and Y) differ greatly in size and structure but have identical DNA sequences within a small localized region. During meiosis I of spermatogenesis, the X and Y chromosomes pair, cross over, and undergo obligatory recombination at that region, ensuring the proper segregation of the sex chromosomes (Raudsepp and Chowdhary, 2015). Because this small sex chromosome region genetically behaves like an autosome (*i.e.*, there are 2 copies in both males and females, and it undergoes recombination), it is referred to as the pseudoautosomal region (PAR).

Although the mouse genome project was completed almost two decades ago, the mouse PAR has remained largely unsequenced, probably because of the high density of repetitive sequences (Takahashi et al., 1994), high GC-content regions (Kasahara et al., 2010), and the absence of bacterial artificial chromosome (BAC) clones containing the PAR sequence of the C57BL/6J (B6J) strain. The latest mouse reference genome, GRCm39, includes only 100–200 kb of sequence within the PAR, punctuated with large gaps.

Assuming there is syntenic conservation with human PAR1 (∼2.7 Mb), comprising 16 genes, a dozen mouse genes are thought to be missing from the mouse genome assembly. The size of mouse PAR in the 129X1/SvJ strain was estimated by physical mapping to be ∼700 kb, the smallest known among mammals (Perry et al., 2001). Because of the obligatory recombination event, the smaller the PAR is, the higher the density (frequency) of recombination will be, and the more likely it is that mutations will occur during meiosis, as recombination involves double-strand breaks. Indeed, the recombination frequency in the mouse PAR is >100 times higher than that in the autosomes and the non-PAR region of the X chromosome (Kipling et al., 1996; Kasahara et al., 2010; Lange et al., 2016).

To date, four genes have been experimentally localized to the mouse PAR: *Mid1*, which spans the pseudoautosomal boundary (PAB) on the X chromosome (Palmer et al., 1997); *Sts*, which was the first identified as a mouse PAR gene (Salido et al., 1996); *Nlgn4*, for which a knockout mouse line was generated before it was identified as a PAR gene (Jamain et al., 2008; Maxeiner et al., 2020); and *Asmt*, which encodes a melatonin synthase that we cloned (Kasahara et al., 2010). The remarkably high mutability of mouse PAR genes was confirmed in *Asmt*, in which a large number of single nucleotide variants (SNVs) and indels were found in many mouse strains. For example, in the B6J strain, two protein-altering SNVs reduce the stability of the ASMT protein and abolish its enzymatic activity. This change deprives B6J mice of the pineal hormone melatonin and, interestingly, confers the evolutionary advantage of accelerated gonadal development in a breeding environment (Kasahara et al., 2010; Zhang et al., 2021). Likewise, the PAR-located portion of the *Mid1* gene appears to have evolved at a considerably faster rate than the X-specific portion (Perry and Ashworth, 1999).

We started with the sequence of a bacterial artificial chromosome (BAC) clone (MSMg01-318O12) containing the *Asmt* gene obtained from a genomic library of the Japanese wild mouse (*M. musculus molossinus*)-derived MSM/Ms (MSM) strain (Abe et al., 2004; Kasahara et al., 2010) and employed PacBio HiFi sequence reads, which were highly accurate and long, to successfully assemble and sequence the entire mouse PAR (Figure 1). Here, we report that the B6J PAR is essentially ∼700 kb, and the MSM PAR is 229 kb; these PAR regions are likely the smallest among all organisms with heterogametic sex chromosomes. This indicates that the mouse PAR has evolved rapidly, and studying its structure at nucleotide resolution provides a blueprint for the PAR and sex chromosomes in general.

**Figure 1.**
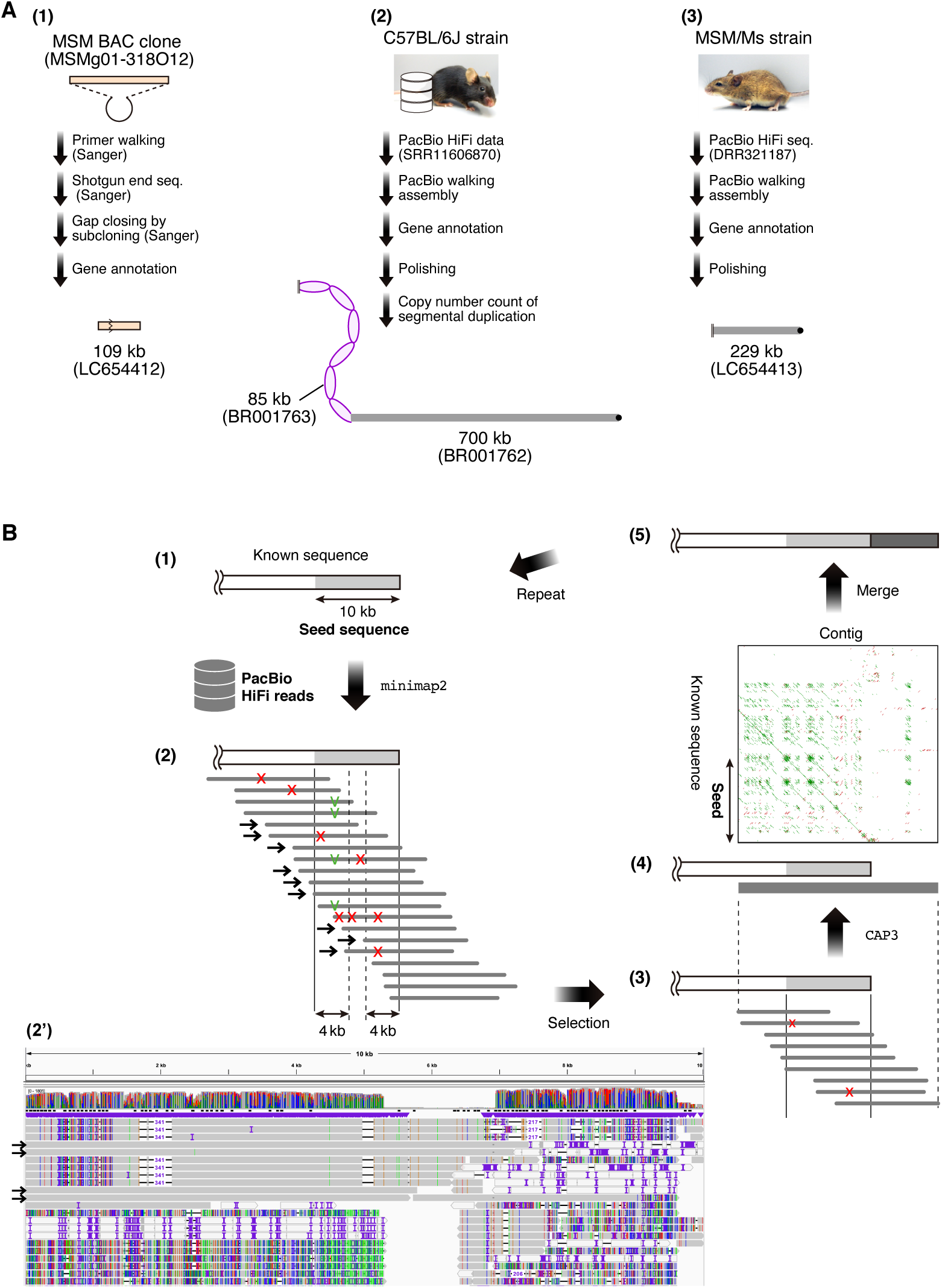
Overview of mouse PAR sequencing. (**A**) Overall workflow for sequencing and assembling the mouse PAR. (**1**) The MSM BAC clone (MSMg01-318O12) containing (a part of) the MSM PAR was sequenced by the Sanger method. The nucleotide sequence (LC654412) is ∼109 kb in length, and the BAC clone harbors a large deletion. (**2**) The B6J PAR was assembled by the PacBio walking method using the publicly available PacBio HiFi read data (SRR11606870). There was a segmental duplication of ∼85 kb, and the copy number was measured by digital PCR. The minimum number of copies was 2, and the average was approximately 6. The PAR sequence containing two copies is deposited in the database as the B6J PAR model sequence (BR001762), and it is ∼700 kb in size. The total size of the PAR containing 6 copies is ∼1 Mb. (**3**) The MSM PAR was assembled by the PacBio walking method using PacBio HiFi sequencing reads from a male MSM mouse. There is no SD in the MSM PAR, and it is ∼229 kb in size (LC654413). (**B**) A schematic diagram of the PacBio walking method. For details, see the Methods section. (**1**→**2**) PacBio HiFi reads showing homology to the terminal 10 kb of a known sequence were extracted using minimap2. (**2**) Red x’s indicate a base (or indel) different from the known sequence; however, since they are found randomly, they represent sequencing errors. Green v’s indicate a base (or indel) different from the known sequence and, since the same variant is found in multiple reads, these changes may represent polymorphisms between repetitive sequence units or heterozygous alleles. An actual image displayed in the IGV is shown in (**2’**). (**2**→**3**→**4**) The reads indicated by arrows were selected and assembled into contigs using CAP3. (**4**) We compared the self-similarity view of the contig with that of the known sequence to confirm that walking proceeded correctly. (**4**→**5**) The contig sequence was merged with the known sequence.

## Materials and Methods

### Animals

MSM/Ms mice (RIKEN BioResource Research Center [BRC] #RBRC00209) maintained in the laboratory animal facility in RIKEN Center for Brain Science (CBS) were used for sequencing and genetic size calculation analyses. C57BL/6J JAX mice were purchased from The Jackson Laboratory Japan. *Mus spretus* (SPR2 strain, #RBRC00208) and *Mus caroli* (Car strain, #RBRC00823) were from RIKEN BRC. *Rattus norvegicus* (F344 strain, NBRP Rat #0140) and *Microtus levis* (MrosA strain) were from Kyoto University and Okayama University of Science, respectively. All procedures were approved by the Institutional Animal Care and Use Committees of the RIKEN, Kyoto University, or Okayama University of Science.

### Immuno-FISH analysis of spermatocytes

Testes from *Mus musculus* (B6J and MSM strains) and *Mus spretus* (SPR2 strain) were detunicated and seminiferous tubules processed according to a previously described protocol for spreading preparations of spermatocytes (Peters et al., 1997). The spermatocyte spread slides were hardened and then denatured for 2 min at 70 °C in 70% formamide/2 × SSC. The BAC clone (MSMg01-38O12) was labeled by nick-translation with biotin-16-dUTP (Cat# 11093070910, Roche Applied Science) following the manufacturer’s instruction. The labeled DNA fragment was ethanol-precipitated with sonicated salmon sperm DNA (Cat# D7656, Merk) and *Escherichia coli* tRNA (Cat# 10109541001, Roche Applied Science), and mouse Cot-1 DNA (Cat# 18440016, Thermo Fisher Scientific) and then denatured at 75 °C for 10 min in 100% formamide. The denatured probe was mixed with an equal volume of hybridization buffer (0.4% BSA, 4 × SSC, 20% dextran sulfate), and pre-incubated at 37 °C for 20 min prior to hybridization. The probe was incubated with the denatured meiotic chromosomes overnight at 37 °C. The spread slides were blocked with undiluted Block Ace (Cat# UK-B80, Dainippon Sumitomo Pharma) for 1 h at room temperature. Immunostaining was performed using a rabbit polyclonal to human SC lateral element protein SCP3 (1:500 dilution; Cat# ab15093, Abcam; RRID:AB_301639) and a Cy3-labeled donkey anti-rabbit IgG (H+L) (1:200 dilution; Cat# 711-165-152, Jackson ImmunoResearch Labs; RRID:AB_2307443). Incubation with primary or secondary antibody was done overnight or for 3 h at room temperature, respectively. Next, to detect the biotin FISH signal, the slides were incubated with FITC-conjugated avidin (Cat# 100205, Roche Applied Science) at a 1:500 dilution in 1% BSA, 4 × SSC for 1 h at 37 °C. Finally, the slides were mounted with VECTASHIELD Mounting Medium with DAPI (Cat# H- 1200, Vector Laboratories). Immunostaining and FISH images were captured using Axioplan 2 imaging microscope (Carl Zeiss) equipped with a CCD camera, CHROMA filter sets, and ISIS4 imaging-processing package (MetaSystems, Germany).

### FISH analysis of metaphase chromosomes

Metaphase slides were prepared from ConA-stimulated spleen lymphocytes or tail-tip fibroblasts from adult males of *Mus musculus* (B6J and MSM strains), *Mus spretus* (SPR2 strain), *Mus caroli* (Car strain), *Rattus norvegicus* (F344 strain), and *Microtus levis* (MrosA strain). The slides were hardened and then denatured for 2 min at 70 °C in 70% formamide in 2× SSC. The denatured, biotin-labeled BAC DNA probe was incubated with the denatured chromosomes overnight at 37 °C. After hybridization, the slides were incubated with FITC-conjugated avidin at a 1:500 dilution in 1% BSA, 4 × SSC for 1 h at 37 °C. Finally, the slides were mounted with VECTASHIELD Mounting Medium with DAPI.

### Sequencing of the BAC clone MSMg01-318O12

The BAC DNA was purified using QIAGEN Large-Construct Kit (Cat# 12462). DNA sequencing by primer walking initially started from both ends of the BAC clone and of every exon of *Mid1*, *Sts*, and *Asmt* genes. Since the mouse PAR is usually GC-rich, we always used a 1:1 mixture of dGTP BigDye Terminator v3.0 (Cat# 4390229, Applied Biosystems) and BigDye Terminator v3.1 Cycle Sequencing kits (Cat# 4337457) supplemented with dimethyl sulfoxide (final 10%(v/v)). DNA sequence data were obtained using ABI3730xl sequencer and were aligned and processed using BLASTN 2 sequences program and a sequence analysis software, GENETYX (Genetyx, Tokyo, Japan).

For shotgun library construction, the BAC DNA was mechanically sheared by HydroShear (Genomic Solutions), and the ends were blunted using Mighty Cloning Kit (Cat# 6026, Takara Bio). DNA fragments of 4–5 kb were fractionated by agarose gel electrophoresis, ligated with pUC118/*Hinc* II BAP (Cat# 3322, Takara Bio), and used to transform *E. coli* JM109. A total of 288 white recombinant colonies (equivalent to ∼12-fold coverage of the BAC clone) were randomly picked, inoculated in 96-well plates, and stored as glycerol stock. The shotgun clones were sequenced from both ends by the Sanger method.

Shotgun clones of informative were sequenced by primer walking. In cases where no appropriate primers could not be designed due to repetitive sequences or sequencing reaction failures, we generated deletion constructs. We purified the plasmid DNA of each shotgun clone was purified using QIAGEN Plasmid Midi Kit (Cat# 12143) and constructed a restriction enzyme map. The insert fragment was extracted and treated with Exonuclease III (Cat# 2170, Takara Bio) typically at 23 °C for 3, 7, 12, 18, 25 min. DNA samples that were shortened to the appropriate size were blunted with S1 nuclease (Cat# 2140, Takara Bio) and T4 DNA polymerase (Cat# 2040, Takara Bio), ligated into pBluescript II KS(+), and sequenced using the Sanger method.

### PacBio walking

A schema of the PacBio walking method is shown in Figure 1B. PacBio HiFi reads (B6J, SRR11606870; MSM, DRR321187) were mapped to known “seed” sequences, *i.e.*, any of the exons of *Asmt*, *Akap17a*, *Mafl-ps*, *Nlgn4*, *Sts*, *Arse*, *Mafl*, *Erdr1*, *Mid1*, and *Gm52481* genes, using minimap2 (ver. 2.17-r941) with the parameters -k 27 -w 18 -m 99. The alignments were processed using samtools (ver. 1.1) and visualized with IGV (ver. 2.8.13). We manually selected reads that were identical (except for obvious mutations or polymorphisms in the seed sequence) to the seed sequence over 4 kb and assemble them using CAP3 (VersionDate, 12/21/07) with the default parameters. If more than two contigs were generated, the contig consisting of the largest number of reads was used as the representative. To visualize the hallmark of the contig sequence, we created a dot-plot view of self-similarity using web YASS (https://bioinfo.lifl.fr/yass/yass.php) or local YASS (ver. 1.15) with the default parameters. We compared the self-similarity view of the contig with that of the seed sequence and confirm that the walking was proceeding correctly. When the self-similarity view of the contig that we took as representative was obviously different from that of the seed sequence, we used another contig as an alternative representative. Next, the representative contig and the seed sequence were compared using BLASTN 2 sequences program (default parameters) and manually merged them. Basically, the contig sequence was connected to the seed sequence near the center where these sequences overlapped. A 10-kb sequence from the end of the merged sequence was used as the new seed for the next round of walking. Each round of walking yielded a new sequence of 3.5–kb (∼9.8 kb on average).

PacBio walking is not a true de novo assembling method, because it needs known correct sequences (>4 kb in this study) and builds new sequences from them. Even HiFi reads of PacBio sequencing contain sequencing errors, but thanks to the known correct sequences, the errors in the mapped and aligned reads can be identified. At the same time, if it is a region of a repetitive sequence containing variants like general minisatellites, it is possible to distinguish between those variants and sequence errors. Heterozygous alleles could also be distinguished. The steps of assembling reads that almost perfectly match a seed sequence to create a contig and merging it into the seed sequence, or the already built sequence, requires considerable care. In most cases, only one contig was generated, but sometimes the CAP3 program generated multiple contigs. To identify the correct contig following the seed sequence, we created and compared dot plots of the seed sequence of the contigs (usually >10 kb each). In other words, we checked the identity of a contig using a dot plot image that characterized the entire sequence, rather than local homology. This is because it is important to correctly merge a correct contig to the seed sequence. The combination of the image characterization of sequences and image recognition technology will make it possible to automate the PacBio walking method, although this will take more computation time than de Bruijn graph-based or overlap-layout-consensus-based assemblers.

### Digital PCR

Genome DNA was extracted from liver, brain, or tail chip using Monarch Genomic DNA Purification Kit (Cat# T3010S, New England Biolabs) and digested by *Pst* I (Cat# R3140S, New England Biolabs). After heat inactivation (60 °C for 15 min) and dilution with water (5 ng/µL), we performed digital PCR using the QuantStudio 3D system (Thermo Fisher Scientific) according to the manufacturer’s procedure. We used TaqMan Copy Number Reference Assay, mouse, Tfrc (Cat# 4458366) to count the chromosome 16 (2 copies per the diploid genome) and Custom TaqMan Copy Number Assays (mMid1Ex5 and mMid1Ex7), which targeted the exons 5 and 7 of *Mid1* gene, designed by the TaqMan Custom Design Assay Tool (Thermo Fisher Scientific). The sequences of the primers and probes are as follows:

mMid1Ex5: CTCAGCAGATTGCAAACTGTAAACA, GGCGTGGTCATTTTCCTTCAG, and [FAM-]TCTGCATCGCTCATCTCGC

mMid1Ex7: CTGCACCGCTTCCTACGA, GTAGGAGACCACGCTGAACTC, and [FAM-]CCACTGGACCTCAGAGGAC

The copy numbers of the SD obtained by using mMid1Ex5 and mMid1Ex7 assays were the same. The copy numbers in DNA samples extracted from the liver, brain, and tail tip of the same individual were the same, indicating that the copy number did not change during ontogeny.

There is a *Pst* I site in the *Mid1* exon 5 of MSM, and a primer and TaqMan probe of the mMid1Ex5 assay contain one base mismatch each to the MSM sequence. So, the mMid1Ex5 assay did not react with the *Mid1* of MSM.

### RT-PCR

Total RNA was extracted from the brain (cerebral cortex), liver, or skin of B6J or MSM using Trizol reagent (Cat# 15596018, Invitrogen) and treated with DNase I (Cat# 2270A, Takara Bio). cDNA was synthesized using SuperScript III First-Strand Synthesis SuperMix (Cat# 18080400, Invitrogen) with oligo(dT) primer or random hexamer. We used LA Taq with GC buffers (Cat# RR02AG, Takara Bio) or Multiplex PCR Plus Kit with Q-solution (Cat# 206152, Qiagen) for PCR and always used a temperature gradient to optimize the annealing. The sequences of the primers are as follows:

*Mafl*/*Mafl-ps*: TCGGGKGACCTCACGCTGAC, GATGACGTCACTTCCTGTTTCC

*Cd99*: TGCAGCCGCGCCCGTCAATCACG, GAGCAGCGTCAGTGATGTCATCG

*Xg*: GGTGATGTCACCCGTGGTCATGG, TCACACCGGCTGCGACTCTGAGG

*Arse*: AGCACGGAACCGGCTGCAGCCTG, CTTCCGGTCCCACAGGAAGTACC

*Akap17a*: ACGCGGAAGAGGAAGTCACCCTC, TTCCTGTCCGTCACTTCCTGTCAC

### Genetic distance calculation

Female B6J mice and male (MSM × B6J)F1 mice were mated to produce a total of 103 progenies (Kasahara et al., 2010) to measure the recombination fractions during male meiosis between the PAB and 17 PAR loci. The genotype of PAB was assumed to be the same as the presence or absence of *Sry* gene. Female (MSM × B6J)F1 mice and male B6J mice were mated to produce a total of 77 progenies to measure the recombination fractions during female meiosis. The genetic distances estimated using Kosambi’s map function.

Genomic DNA was extracted from mouse tail biopsies by proteinase K–SDS digestion. Genotyping of the loci was performed by PCR and direct sequencing of the PCR product or size estimation on an agarose gel. The primers used are as follows.

*Sry* gene: GGAGGGCTAAAGTGTCACAGAG, TGGTTTTTGGAGTACAGGTGTG

DXMit81: TGGCAGCACTTTAAGCATTG, TTCCCAAGCTGCTGTTTCTT

DXMit25: GAGGTGGGGAGAAACAGAGG, GAGGAGCATCAACCTTCTCG

DXMit97: AAAATGGATGGGGAAAGGAG, CATTTGAAGTCAATATCAGTCATCG

DXMit249: AAAATAGAACTTCAGCAGCATGC, TTATGTGCTTATTAGCCAAGGTG

*Mid1* (Intron 2): ATGCTCAGCCTGAAAATCAAG, TCGCCCAACTCTCTCACTATGTT

*Mid1* (Exon 5): AGGCTCCGCAARTTAGCTCAGC, CAGTGATATTCTTTGCKGTCTG

*Mid1* (Exon 7): AGAGAAGAGCTCTGCACCGC, ACATTGGCYTGTCCGGTGAA

*Mid1* (Exon 8): CTGTGTAACTCGGCGGACAG, TCACCGTGAAGATRTACTTGG

*Mid1* (Exon 10): ATCGGCCTGGCGTACAGATC, GGTCAGACACTTGTTCCACACG

*Mafl* (Intron 5–Exon 6): CCTGTCTGGTGGCCATGAC, CGGCGAGAACGTCACGTG

*Mafl* (Intron 1): AAGGAGATCCTGGGGCTCAC, CAGCCTGTCACTCATACTTA

*Cd99* (Intron 7): AGGTAGGRGACAGGAAGTCC, GGACAGGAAGTGACGTCACG

*Xg* (Intron 3): ATCCACGCCCAAGCCAGGG, TGGGGCAGCCGCGGGTAGAG

*Arse* (Exon 9–Exon 10): GATGTCCTGCTCCACTACTG, GCGTCACGAAGTCAATTTCCAC

*Arse* (Exon 4–Exon 5): CCCAGGAGCTGACATTCG, GTCTCGCAGTTCAGGCCAAG

*Sts* (Exon 10): TGCAGCCACGGCTTCCAC, CAGCGCRGTGTGGAAGGC

*Sts* (Exon 2–Intron 2): GCTCGCTGACATCATCCTC, GACGCCGTGAACAGGTACACG

*Nlgn4* (Exon 6): GCTCCCAAAGGTGTTGAACC, TGCTCTTCCTCAACGTCCTC

*Nlgn4* (Intron 3): AGCCGGTCATGGTTTACATC, GAGTAGTGCGACAGCGTGAG

*Mafl-ps* (Exon 5): CGGATGTCGCTGACGTCACC, GCGGCCTTGCTCAGCGTCTC

*Akap17a* (Exon 1): ATCTCGAACTGGGAGGTGATG, GAAGCTGAAGGTGTGGAAGC

*Asmt* (Exon 6): CAAGCCCTCAGGGTTCAGGAA, GCAAAGTGAACGCCGATGTG, CGAGGTCACCGTGTTCGAGA (for sequencing)

#### Minigene splicing assay

The full-length of mouse *Asmt* gene (MSM) along with 5’- and 3’-UTR sequences was obtained by PCR using the BAC clone DNA as a template and the following primers: AGGCTCAGTATCTCGCGTCCCACGATG and CGCAGACACCAGATGGCTGTAACTGAC. The PCR product was inserted into pCR2.1 vector (Cat# 451641, Invitrogen), verified by sequencing, and then subcloned into the blunt-ended *Apa* I site of pcDNA5-FRT-TO (Cat# V652020, Invitrogen).

The human *Asmt* exon 4, 5, or 6 along with its flanking intron regions containing appropriate restriction enzyme sites was synthesized (Biomatik or Genewiz). Each exon of mouse *Asmt* gene in pcDNA5-FRT-TO was replaced with the human form using the appropriate restriction enzymes. The constructs were integrated into a specific genomic location of Flp-In T-REx 293 cells (Cat# R78007, Invitrogen) and stable cell lines were selected according to the manufacturer’s instructions.

Induction of the expression of *Asmt* by the addition of doxycycline (0.5 µg/mL; Cat# 631311, Takara Bio) for 24 h and total RNA was extracted using Trizol reagent. One stable cell line was used for each minigene construct, and RNA was collected after the expression induction only once. After DNase I treatment, cDNA was synthesized using SuperScript III First-Strand Synthesis SuperMix and BGHpA-Rv primer (CAACTAGAAGGCACAGTCGAGG) targeting the 3’-UTR (polyadenylation signal) sequence of pcDNA5-FRT-TO vector. cDNA was amplified with *LA Taq* DNA polymerase using the primers mAsmt-Ex3-Fw (CCAACTCCCCCCTGGCGTCCAC) and BGHpA-Rv. RT-PCR products were cloned into pCR2.1 vector and used to transform *E. coli* TOP10 F’ (Cat# C303003, Invitrogen). Twenty white colonies were randomly picked up for each construct/cell line and sequenced.

## Results

### Sequencing of the mouse PAR (C57BL/6J and MSM/Ms)

Since the end sequences of the BAC clone (MSMg01-318O12) were localized to *Mid1* intron 5 and the subtelomeric region (Kasahara et al., 2010), we initially hypothesized that the BAC clone would contain the majority of the mouse PAR. Fluorescence *in situ* hybridization (FISH) analysis using the clone in meiotic pachytene spermatocytes confirmed that it localized to a synapsed region of the sex chromosomes in mice (both B6J and MSM strains) (Figure S1A, B). We first sequenced the BAC clone using the primer walking method and then by constructing a shotgun library. We closed sequencing gaps by subcloning informative shotgun clones and obtained the entire sequence of the BAC clone. We identified the *Mid1* (exons 6–10), *Sts*, *Nlgn4*, *Mafl-ps*, *Akap17a*, and *Asmt* genes (Figure S2A, B); however, the BAC clone was considered insufficient to provide a complete structure of the mouse PAR because the insert size was 109 kb, which was inconsistent with the previously estimated PAR size (∼700 kb) (Perry et al., 2001). In addition, two candidate PAR genes, *Arse* (discovered as mouse expressed sequence tags (ESTs) by sequence homology with human *ARSE*) and *Mafl* (identified during RT–PCR analysis of *Mafl-ps*), were missing. We now understand that the BAC clone contains an ∼93-kb deletion (Figure S2).

A PacBio HiFi sequencing dataset of the mouse genome (B6J) has been published recently (Hon et al., 2020). Although a BLASTN search revealed that several reads contained *Sts*, *Asmt*, or *Arse* sequences, we could not obtain a faithful assembly of the PAR using typical *de novo* assemblers, possibly due to the presence of massive repetitive sequences. The analysis of the BAC clone revealed that unique alignments were possible with sequences of >4 kb even in repetitive regions, and we ascertained a rough map of the mouse PAR (*Mid1*–*Mafl*–*Arse*–*Sts*–*Asmt*–telomere). Thus, we hypothesized that it would be possible to order and connect the PacBio HiFi reads using a primer walking-like method (Figure 1B).

The “PacBio walking” method enabled the assembly of highly problematic sequences, such as a large repetitive region (∼130 kb) and ∼5,400 repetitions of a 25-base consensus sequence (see Figure 5), allowing us to obtain a draft sequence of the entire PAR of the B6J strain (Figure 2A). The B6J PAR comprises 10 genes and 4 pseudogenes and is a minimum of ∼700 kb in size, depending on the copy-number polymorphism of a segmental duplication (SD) described later (Figure 2B).

**Figure 2.**
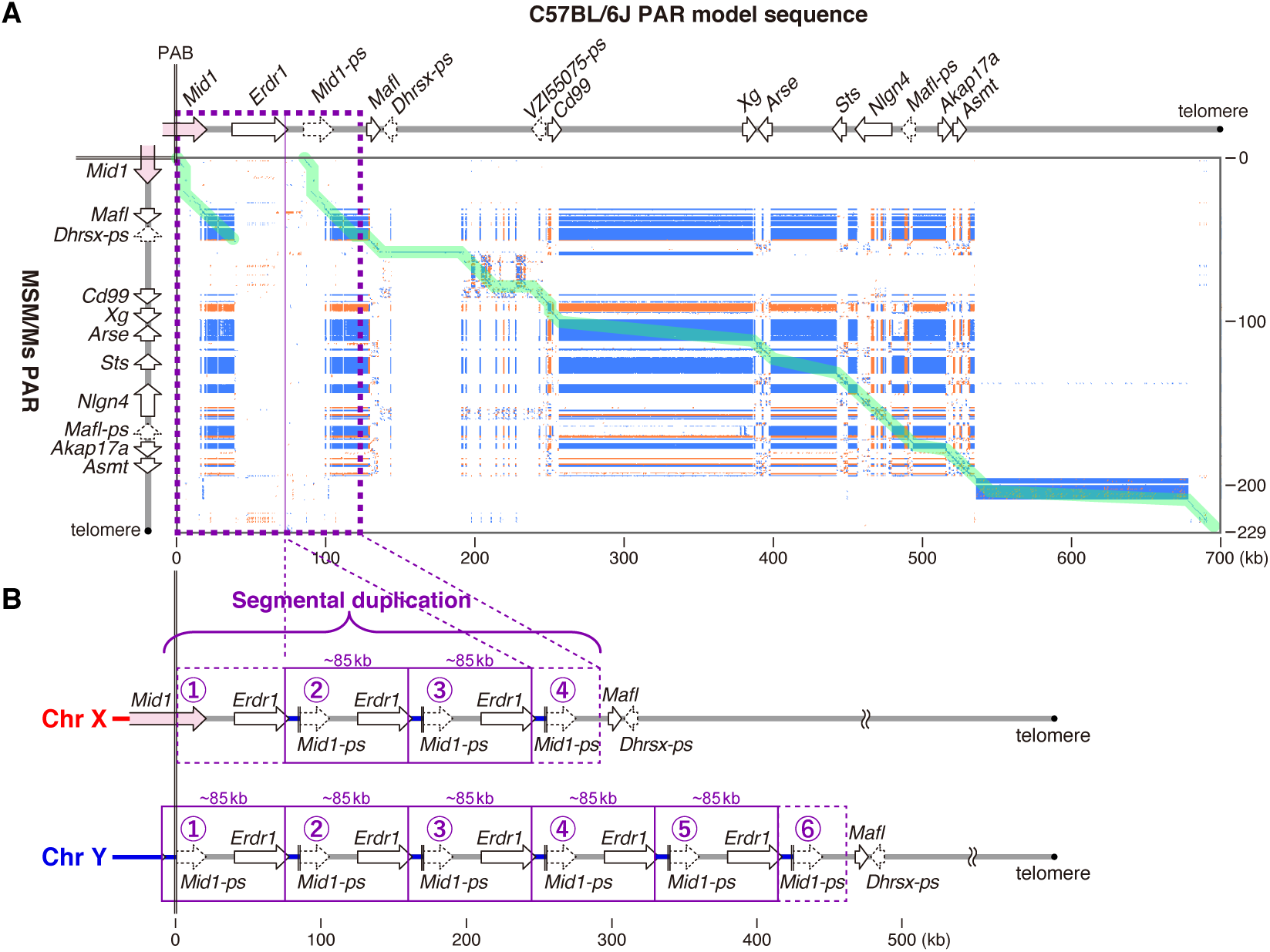
Structure of the mouse PAR. (**A**) A dot plot showing similarity between the B6J and MSM PAR sequences. The B6J PAR model sequence, including only the first and last segmental duplication (SD) units, contains 10 genes and 4 pseudogenes (the ’-ps’ suffix denotes pseudogene). Forward and reverse complement matches are shown as blue and orange dots. Green lines indicate a possible alignment between two sequences. The SD region present in the B6J PAR is indicated by a dashed rectangle. (**B**) The SD structure. The SD has CNVs (2 to >10), and as an example, XY chromosomes with 4 and 6 copies of SD are shown. Each SD unit (85.5 kb) contains 8.5 kb of a Y-derived stretch (which includes *Erdr1* exon 3), the PAB, *Mid1* exons 4–10 (*i.e.*, *Mid1-ps*), and *Erdr1* exons 1–2. The SD copy number is defined by the number of *Mid1*/*Mid1-ps*.

The MSM PAR sequence was also determined by the PacBio walking method using a PacBio HiFi sequencing dataset of genomic DNA extracted from a single male MSM mouse. The entire length of the MSM PAR is 229 kb (Figures 2A and S2), which is much smaller than the B6J PAR; however, the MSM PAR contains all the PAR genes except *Erdr1,* which is lacking as the MSM PAR has no SD structure. This smaller-sized MSM PAR with fewer repetitive sequences is consistent with a previous observation made by Southern blot analysis (Takahashi et al., 1994).

### Accuracy estimation of the mouse PAR sequences

We attempted to polish the draft sequences by using standard programs with long and/or short reads. However, this decreased the accuracy in and around repeated sequences. Thus, we focused on the coding sequences (CDS) of the PAR genes and a transcribed pseudogene, polishing the sequence information using sequence data from EST clones and RT–PCR. All errors detected were single-nucleotide indels in long homopolymers, and the error rate of the draft sequences was 1 error/∼3.5 kb or lower.

A comparison of the sequences of the MSM PAR and the original BAC clone might have allowed an estimation of the accuracy of the draft sequence (Figure S2); however, there were so many differences/variants that such an estimation was not possible. It is currently unclear whether these variants arise from errors in PacBio HiFi sequencing/assembling, instability of the BAC clone, errors in Sanger sequencing, or the evolution of the MSM PAR during the ∼20 years between the construction of the BAC and the PacBio libraries.

### Segmental duplication in the B6J PAR

We discovered that there are ∼85.5-kb SDs with high sequence identity that cannot be distinguished by the PacBio walking method (Figure 2B). The first unit of the SD starts ∼8.5 kb proximal to the PAB in the Y-unique region and ends in *Erdr1* intron 2. The first SD unit on the X chromosome is an incomplete structure ∼8.5 kb shorter. The most distal, final unit is also an incomplete structure that does not contain exons 1 and 2 of *Erdr1*, and *Mafl* is located just distal to this structure. We define the number of *Mid1*/*Mid1- ps* copies as the copy number of the SDs (Figure 2B).

Illumina short-read data from one B6J female mouse (Sarsani et al., 2019) confirmed that SD was confined to a region (PAB–*Erdr1*–*Mid1-ps*) in the B6J PAR (Figure 3A), but it was difficult to determine the copy number. We previously observed a polymorphic variation in length around the *Mid1* gene among B6J animals by Southern blot analysis (Kipling et al., 1996; Palmer et al., 1997). Recently, Acquaviva and colleagues also observed a polymorphic variation in PAR length by immuno-FISH imaging (Acquaviva et al., 2020). These observations indicate that the polymorphism might be due to SD copy-number variation (CNV). We estimated the copy number by quantitative digital PCR targeting *Mid1* exon 5 of B6J and found that it ranged from 4 to 25 per diploid B6J genome (Figure 3B). Each generation resulted in new diverse CNVs, and as a result, individual identification by DNA analysis is possible even in the inbred B6J strain. We examined F1 hybrid mice from crosses between B6J and MSM using the digital PCR method that did not detect MSM DNA, confirming that CNV variation arises during spermatogenesis in the B6J mice, not during oogenesis (Figure 3B).

**Figure 3.**
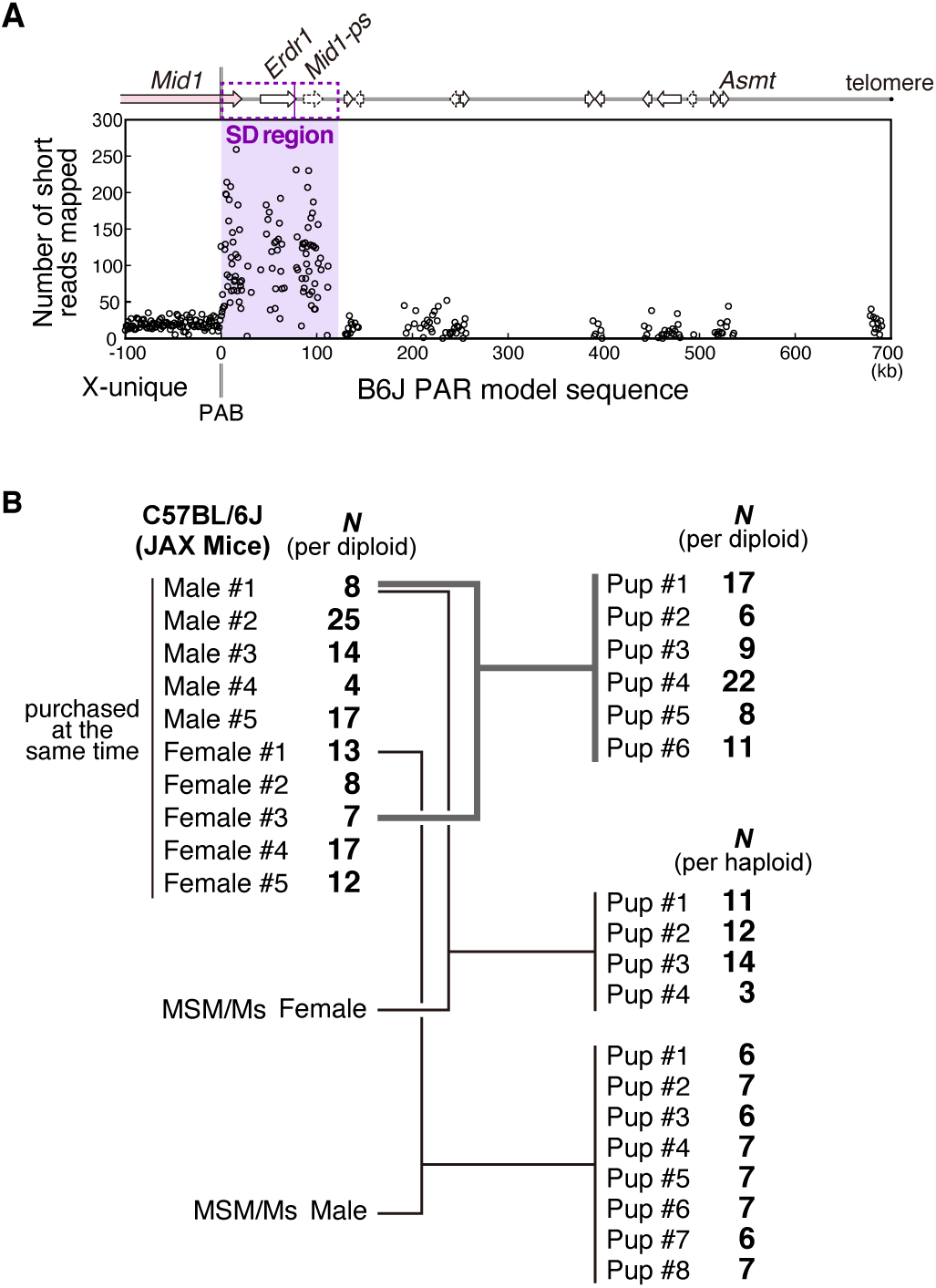
Estimation of the SD copy number. (**A**) An estimation based on numbers of short reads mapped to the PAR. We counted the short reads (from one female mouse) that exactly matched 200-bp sequences approximately every 500 bp, excluding repetitive sequences. The median numbers of mapped reads to the X-unique, the SD, and the other PAR regions were 19, 99, and 13, respectively. (**B**) Estimation of the copy number by digital PCR targeting *Mid1* exon 5. The copy numbers (*N* per diploid or haploid genome) in 10 purchased B6J JAX mice, 6 pups of the next generation, 3 F1 pups (MSM x B6J), and 8 F1 pups (B6J x MSM) were rounded to integer values. The raw measurements are as follows: 8.22, 24.63, 14.10, 3.86, 16.81, 12.73, 8.40, 6.96, 16.99, 12.29; 17.06, 6.04, 8.89, 22.07, 7.87, 10.76; 10.70, 12.00, 14.11, 3.24; 6.11, 7.46, 6.03, 7.01, 6.83, 7.30, 6.25, 7.21. Note that this digital PCR did not react with *Mid1* in the MSM strain.

We created a model sequence of the B6J PAR (∼700 kb) containing only the first and last SD units (*i.e.*, a copy number of 2 per haploid) (Figure 2A) and used this sequence for subsequent analyses.

### Identification and cataloging of the mouse PAR genes

We predicted protein-coding genes in the mouse PAR using the AUGUSTUS, FGENESH, and GeneScan programs and checked mRNA expression of the predicted candidates by searching the EST database and mapping public RNA-seq data to the B6J PAR sequence (Söllner et al., 2017) (Figure S3). We identified 10 genes, 1 transcribed pseudogene and 3 untranscribed pseudogenes in the B6J PAR, and comprehensive information on these genes (the mouse PAR gene catalogue) is provided in Supplementary Note. Many of the genes present in the PAR of humans and other eutherians have been transferred to autosomes or the non-PAR of the X chromosome in mice or lost entirely (Figure 4).

**Figure 4.**
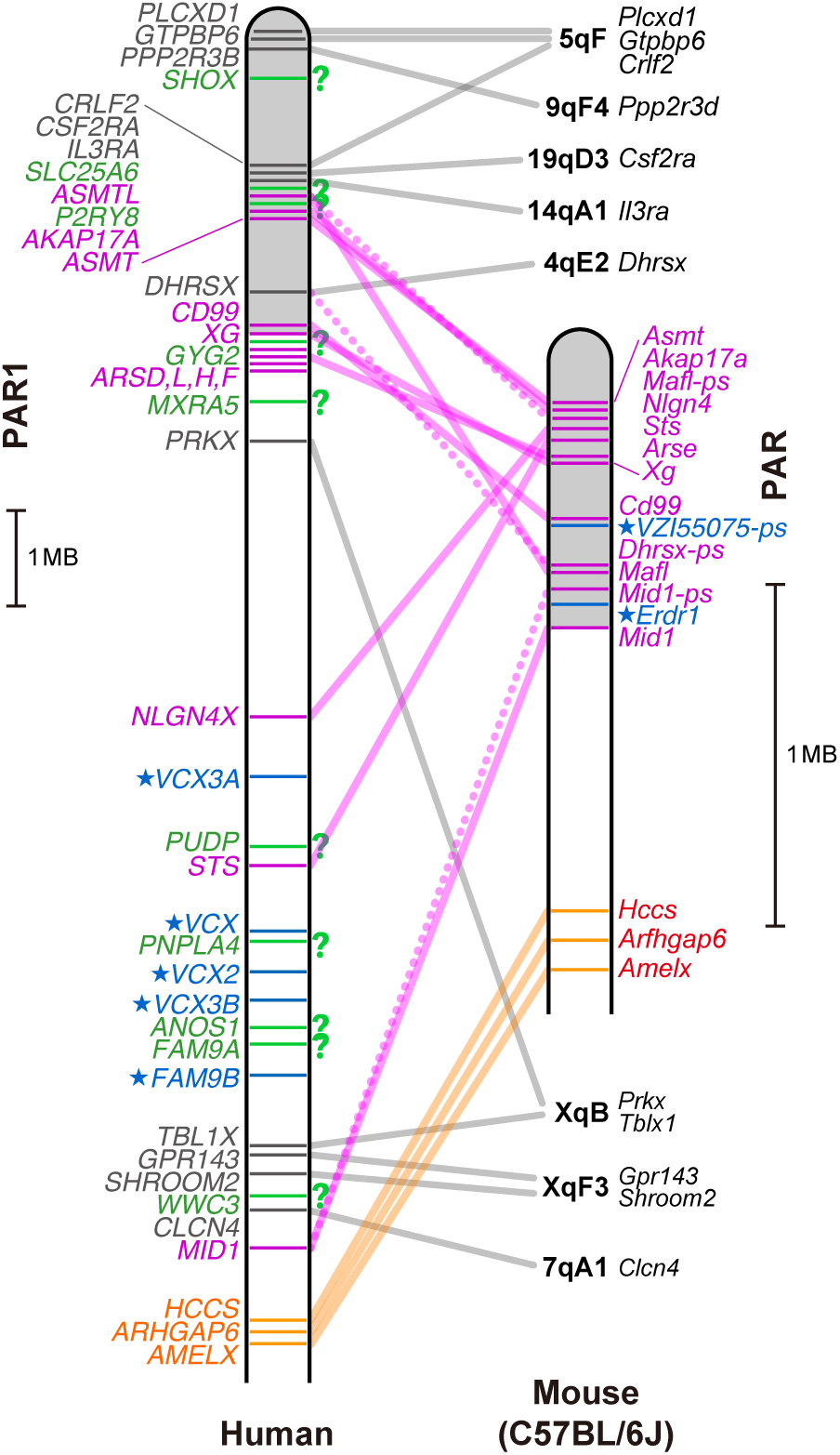
Comparison of mouse PAR genes and human PAR1 genes. Orthologous pairs of human genes located in the PAR1 or the PAB region of the X chromosome and mouse genes are connected by lines. *AMELX* is thought to be located at the PAB in the ancestors of eutherian mammals (Katsura et al., 2012). Mouse PAR genes are shown by magenta connecting lines. Mouse pseudogenes are connected by dotted lines. Stars indicate genes unique to mice or humans. Question marks indicate human genes for which no orthologs have been found in the mouse genome. Orange lines connect human and mouse genes that are both in the PAB region of the X chromosome. Gray lines connect pairs of human PAR1/PAB genes and mouse genes that have moved to autosomes or other regions of the X chromosome.

### Massive mo-2 minisatellites in the mouse PAR

The mouse PAR sequence is characterized by massive repetitive structures, most of which are mo-2 minisatellites consisting of tandem repeats of a 31-bp consensus sequence (Takahashi et al., 1994; Pardo-Manuel de Villena and Sapienza, 1996) or mo-2- like sequences (Figure 5). These minisatellites account for 22.6% of the B6J PAR. Although the mo-2 minisatellite is involved in the structure and behavior of the PAR sister chromatids during mouse male meiosis (Acquaviva et al., 2020), it is unknown whether mo- 2-like minisatellites function in the same way.

**Figure 5.**
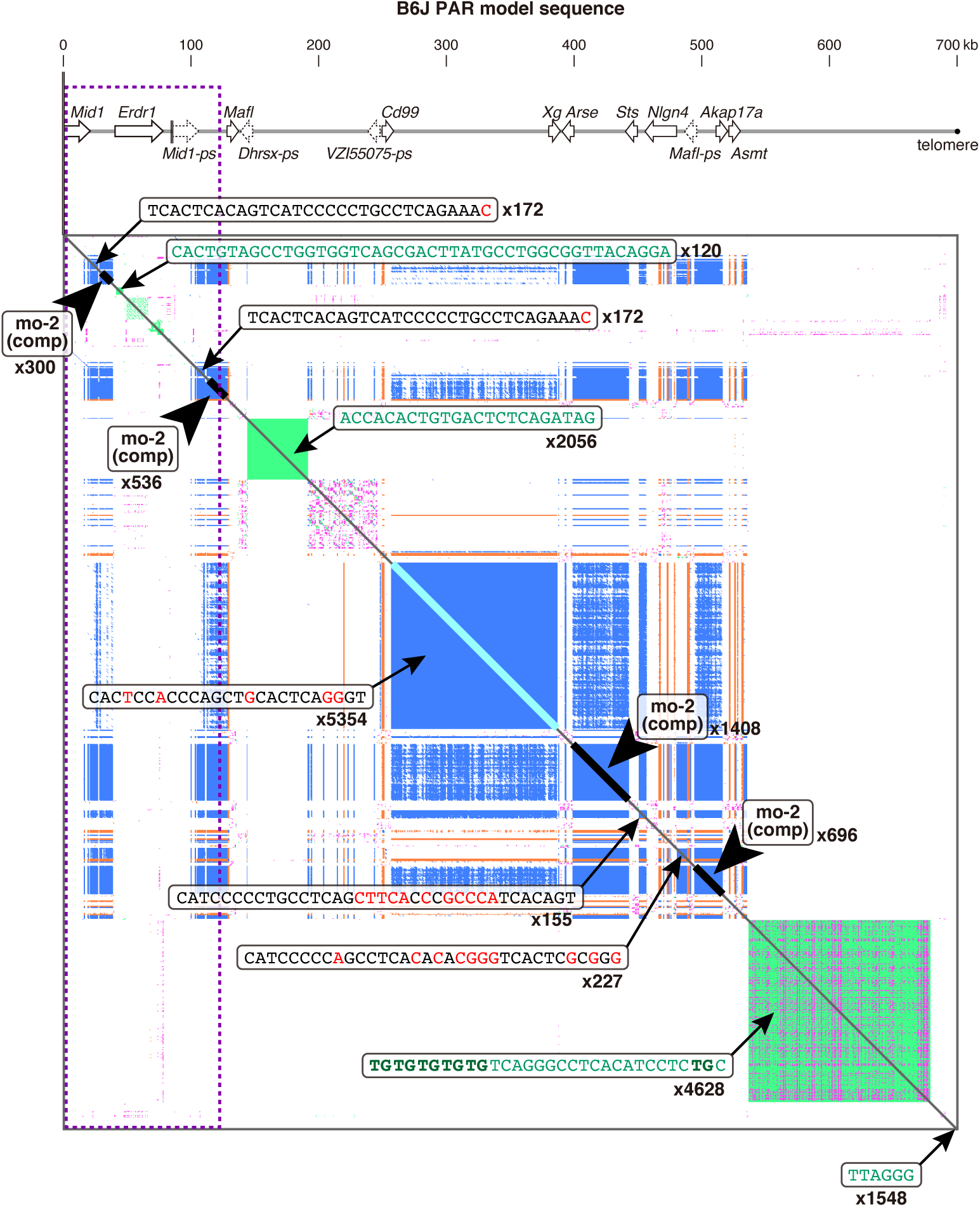
Large minisatellites in the mouse PAR. A dot plot of the self-similarity of the B6J PAR model sequence. Minisatellites consisting of the mo-2 sequence or the complementary mo-2 sequence are shown in blue and orange, respectively. Minisatellites shown in green or magenta have a different sequence from mo-2. The consensus sequences and the repeat numbers of larger minisatellites are shown on the dot plot. Minisatellites labeled “mo-2 (comp)” have a consensus sequence (TCACTCACAGTCATCCCCCTGCCTCAGAAAA) that is the complementary sequence of mo-2. Consensus sequences similar to mo-2 are shown in black letters, and nucleotides mismatched from mo-2 are shown in red letters. Consensus sequences of minisatellites that are not related to mo-2 (including the telomere sequences) are shown in green letters.

The GC content bias of the mo-2 sequence is not particularly high (52%) but is still greater than that of the non-PAR X chromosome (39%) (Figure S4) and the genome-wide average (41%). Since the mo-2 sequence contains more purine nucleotides (heavier nucleotides) than pyrimidine nucleotides and mo-2 complementary sequences predominate in the PAR, the PAR-wide purine content (B6J model, 44.4%) is lower than the genome-wide average (50.0%) and that of the non-PAR X chromosome (50.0%) (Figure S4). Thus, the plus (+) strand (encoding *Mid1*) is lighter than the minus (-) strand, much like the relationship between heavy and light strands of mitochondrial DNA. The strand compositional asymmetry and the large variation in weight depending on the nucleotide position may affect the physical behavior of the mouse PAR.

### The genetic size of the mouse PAR

We investigated genetic recombination in the mouse PAR during meiosis in F1 mice crossed with B6J and MSM by genotyping 17 loci in the PAR (Figure 6A). The genetic distance between the PAB and *Asmt* exon 6, the most distal locus genotyped, was 54.4 cM during male meiosis (Figure 6B). The recombination frequency was approximately 63 or 290 cM/Mb depending on whether the sequence length of B6J or MSM was used, respectively, ∼100- to ∼500-fold higher than the genome-wide average of mouse chromosomes (Jensen-Seaman, 2004) and tens of times higher than that of human PAR1 (Flaquer et al., 2008).

**Figure 6.**
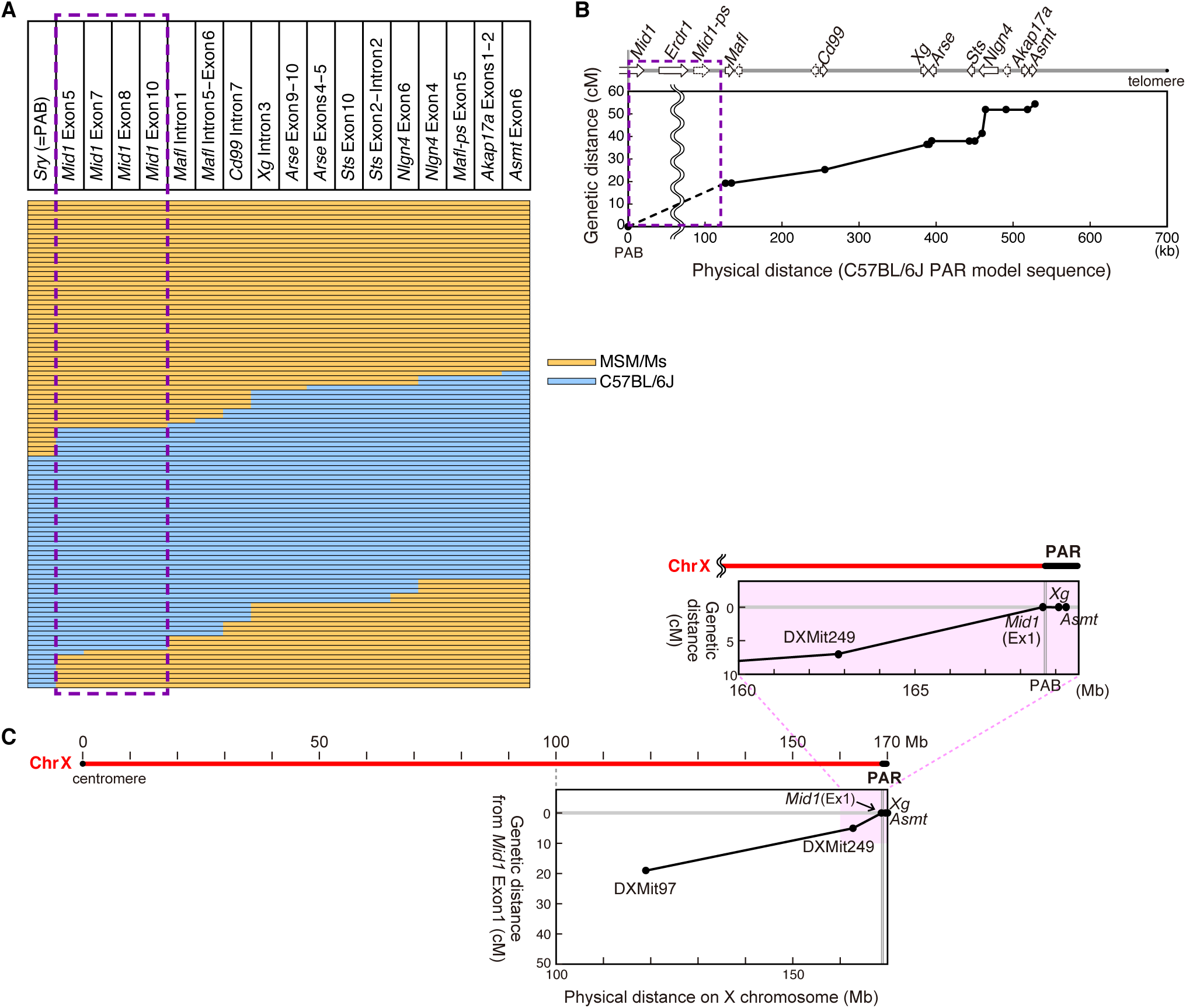
Recombination in the mouse PAR during meiosis. **(A)** Mouse PAR haplotype map of meiotic products in male (B6J x MSM)F1 mice (*n* = 103). The yellow and cyan bars represent the sequences (alleles) of the MSM and B6J strains, respectively. The SD region is indicated by a dashed rectangle. SNVs in the exons of the *Mid1* gene were assayed by PCR–direct sequencing to genotype the 4 loci in the SD region. Therefore, in some cases, a single copy of MSM-derived alleles was sufficient for detection, while in other cases, more than half of the SD copies would have to be MSM- derived alleles. (**B**) Genetic map distance of the mouse PAR during male meiosis. Genetic distance was plotted against the B6J PAR model sequence. The SD region present in the B6J PAR is indicated by a dashed rectangle, and the genetic distance data cannot be plotted in this region. (**C**) No recombination occurred in the mouse PAR during female meiosis (*n* = 77). The loci of *Asmt*, *Xg*, *Mid1* exon 1 (X-unique, ChrX: 168.5M), DXMit249 (ChrX:162.8M), and DXMit97 (ChrX:118.3M) were genotyped.

The reason for recombination in the small region of PAR has been thought to be an obligatory crossover (Burgoyne, 1982; Flaquer et al., 2008). Although we detected a recombination event in only ∼40% of PARs during meiosis in the spermatocytes of F1 mice (Figure 6A), it should be noted that, in B6J males, an unequal crossover event almost always occurs in the SD (Figure 3B), and additional recombination may occur in the remaining PAR. We also examined recombination events during female meiosis (*n* = 77), but no recombination was observed in the small PAR (Figure 6C), as expected due to recombination occurring throughout the X chromosome in females.

### Three major characteristics of mouse PAR genes

There are three characteristics common to the mouse PAR genes that have been rapidly evolving: (i) high-density repetitive sequences in introns, (ii) short introns, and (iii) extremely high GC content in exons. These were first suggested by comparing mouse *Mid1* exons 1–3 located outside the PAR and exons 4–10 located in the PAR (Perry and Ashworth, 1999) and by the *Asmt* sequence (Kasahara et al., 2010). Complete sequencing revealed that these characteristics are true for all mouse PAR genes except *Erdr1*. In addition, genes that existed in the ancestral PAR have moved to autosomes in the rodent lineage; thus, mice have retained these features (Table 1).

(i) Most PAR gene introns are occupied by various types of repetitive sequences, including mo-2 and mo-2-like minisatellites (Figures 5 and S5B). These high-density repetitive sequences likely arise due to high-frequency recombination and unequal crossover events.
(ii) Mouse PAR genes are shorter in length than their human orthologs (Table 1). The length of the CDSs (exons) is almost the same as that of the human orthologs; thus, the shorter length of the PAR genes is due to the short introns.
(iii) The GC content of mouse PAR gene exons is much higher than that in introns and intergenic regions (Figures S4 and S5A). In particular, the GC content at third codon positions (GC3) is extremely high, much higher than the average for all mouse genes (58.7%), with the highest being 96.7% in *Nlgn4* (Table 1). The basic amino acid residues lysine and arginine are of interest. Not only was the third position of the codon replaced with G or C, but most lysine codons (AAA, AAG) were also replaced with arginine codons (CGC, CGG), maximizing GC content. We introduce the RK value as the ratio of the number of arginine residues to total basic amino acid residues (R/(K+R)) for each gene. The RK value of every PAR gene product was higher than the average for all mouse proteins (50.5%), with the highest being 100% in *Asmt* (Table 1).

**Table 1.**
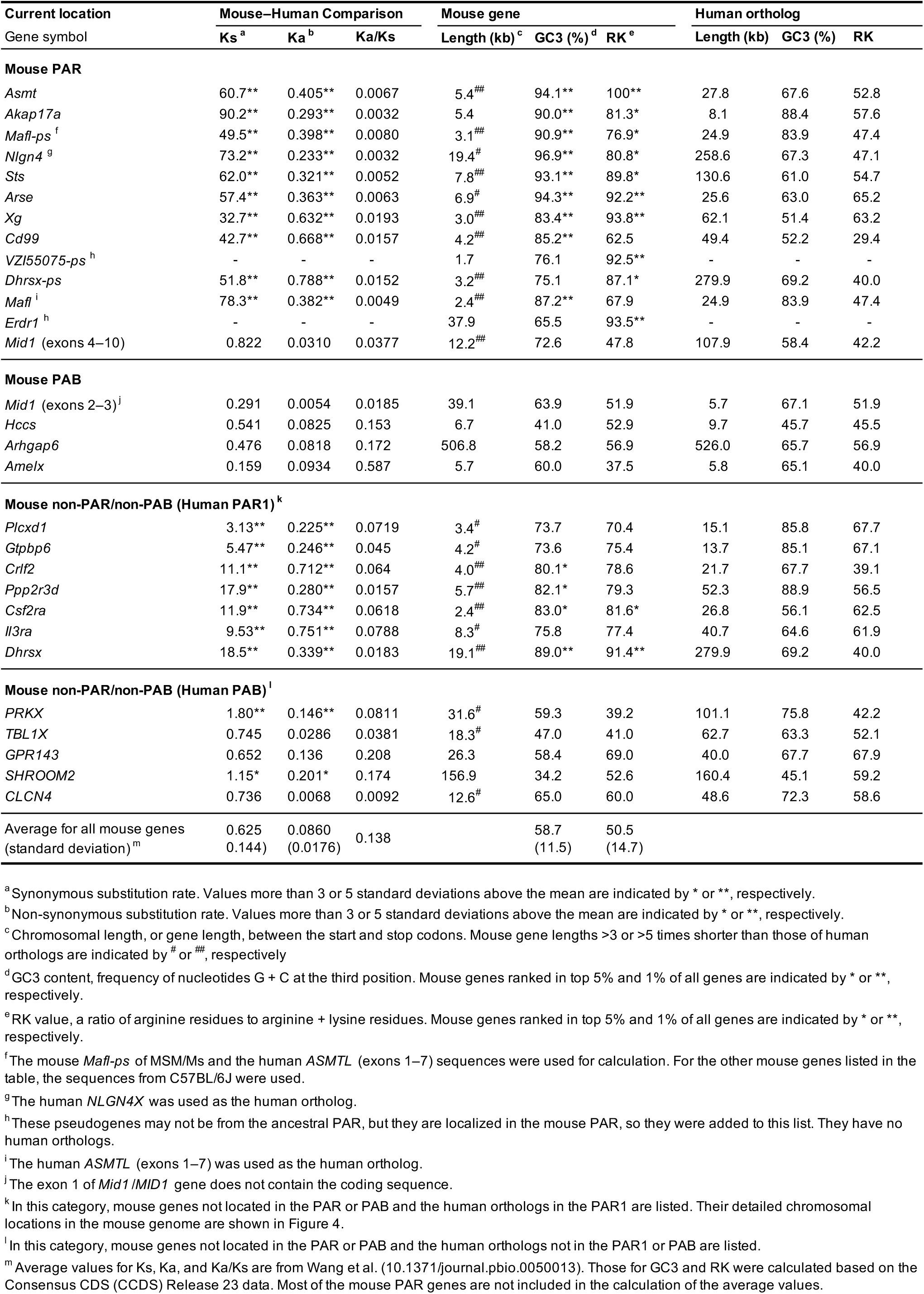
Charateristics of mouse genes whose orthologs were in the ancestral PAR

Exons and splice sites generally have a higher GC content than flanking introns, which may be involved in exon recognition by the spliceosome (Amit et al., 2012). We examined the relationship between the higher GC content of exons and splicing efficiency using mouse *Asmt* as a significantly evolving gene. A minigene in which exons 4, 5, or 6 or both exons 4 and 6 of the mouse *Asmt* gene were replaced with corresponding exon(s) of human *ASMT*, which had a lower GC content (Figure S5A), was recombined into HEK293FT cells to analyze the structure of the mRNA expressed from the minigene. mRNAs with abnormal splicing were significantly more abundant (Figure S5C, D).

Considering that short introns are also involved in the recognition efficiency of exons by the spliceosome (Roy et al., 2008), the high GC-content exons and short introns are likely to be the consequence of a high recombination frequency, leading to the proliferation of minisatellites in introns and intergenic regions. Potentially, this is a reinforcing feedback loop in which an increase in the recombination rate shortens introns (and the PAR itself), further increasing in the recombination rate.

## Discussion

In this study, we developed the PacBio walking method that robustly resolved repetitive sequences and revealed the entire mouse PAR sequence by using highly accurate long-read sequencing data. Elucidating the PAR sequence completes the euchromatic genome sequencing of the mouse.

Sequencing the mouse PAR confirmed that it is indeed small; however, small PARs are not limited to mice and may be common across rodents. This is supported by the high GC3, high RK value, and short length, which are related to each other, of the *Asmt* gene in Rodentia (rodents) or Glires (rodents, rabbits, pikas) (Figure 7). The fact that the X and Y chromosomes do not form synapses or chiasmas (or do not pair tightly) during meiosis in several rodents (de la Fuente et al., 2007; Borodin et al., 2012; Dumont et al., 2018) also supports this assumption. The rat sex chromosomes are also asynaptic (Joseph and Chandley, 1984), and the rat genome sequence shows that many of the original PAR genes are located on autosomes or the non-subtelomeric region of the X chromosome. For example, rat *Asmt* is located on chromosome 12 and has a high RK value and short gene length, while GC3 is exceptionally low (Figure 7A), suggesting that the ancestor of rats had a short PAR and that the PAR was subsequently lost in rats.

**Figure 7.**
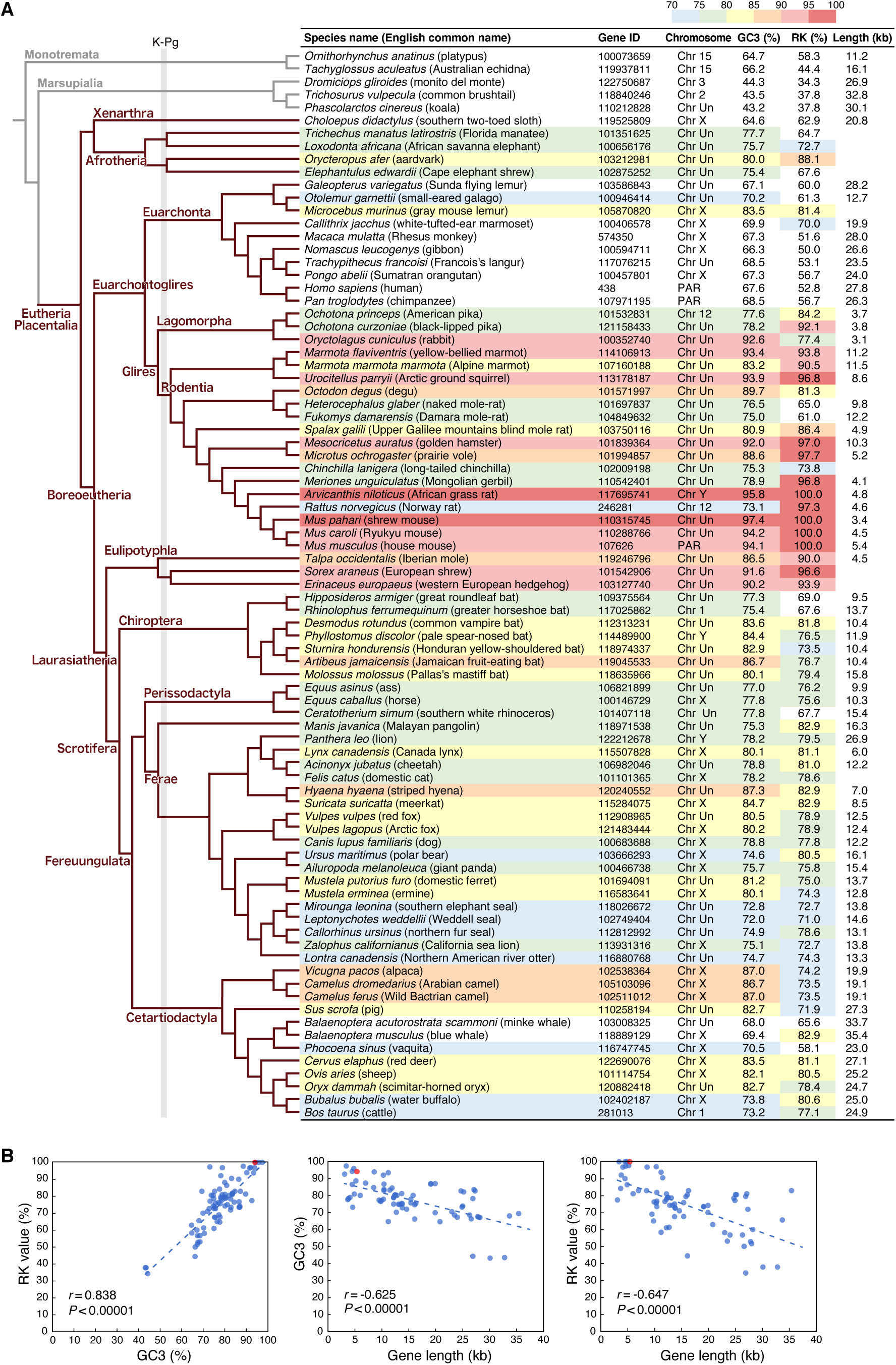
GC3, RK value, and gene length of the *Asmt* gene in 83 mammalian species. (**A**) A list of mammalian *Asmt* genes. The GC3 and RK values of *Asmt* are very high in not only Rodentia but also Eulipotyphla species (hedgehogs, shrews, and moles), suggesting that they have (or had) small PARs. Indeed, many species of *Eulipotyphla* also have asynaptic sex chromosomes, transformed sex chromosomes, or a novel sex-determination system. K-Pg, Cretaceous–Paleogene boundary. (**B**) Relationships between GC3, RK value, and gene length of the mammalian *Asmt* genes. The correlation coefficient (*r*), *P* values, and regression lines are shown. The mouse *Asmt* gene is indicated by a red dot.

The rat has obtained a PAR-independent sex chromosome segregation system, in which they associate in opposite directions during male meiosis (Matveevsky et al., 2021). On the other hand, mice have acquired mo-2 arrays, likely ensuring proper segregation of the sex chromosomes. FISH analysis using the MSM BAC clone, of which 25.3% of the total length is the mo-2 sequence, demonstrated that the mo-2 arrays appear to be present only in mice and their closest relative, *M. spretus;* the next closely related species, *M. caroli*, does not possess mo-2 arrays (Figure S1). The mo-2 array was not detected in rats or *Microtus levis,* which possesses transformed, large-sized sex chromosomes (Borodin et al., 2004). Thus, rodent species may each have unique methods for circumventing male meiosis issues resulting from small PARs; additionally, some species have lost the Y chromosome (Honda et al., 1977; Fredga, 1988; Matveevsky et al., 2017), and segregation is not a concern. Meiotic failure during meiosis in spermatogenesis causes male infertility and creates a reproductive barrier (Schilthuizen et al., 2011). Therefore, the small PAR could readily lead to a reproductive barrier and speciation and explain why rodents are the order of mammals with the most species.

Obtaining the complete sequence of the mouse PAR paves the way for exploring the structure and epigenetics at nucleotide resolution, which will provide a blueprint for the PAR and sex chromosomes. Since the mouse PAR has already changed in sequence, size, and arrangement between strains, mice will likely further speciate into several distinct species in the near future, and the very small PARs of the neo-mice are likely to be disrupted and inefficient in the pairing process. Therefore, these neo-mice will need to acquire a new mechanism to ensure meiotic segregation of sex chromosomes, as found in other current species, or achieve a new sex-determining system that lacks a Y chromosome; otherwise, these species will lose fitness and may become extinct. Sequencing additional Rodentia and Eulipotyphla genomes (Figure 7A) will allow a more reliable prediction of the future evolution of PARs and sex chromosomes.

## Acknowledgments

We are grateful to the Support Unit for Bio-material Analysis (RIKEN). This work was supported by RIKEN President’s Discretionary Fund (to T.Kas.), RIKEN Incentive Research Grant (to T.Kas.), grants for Career Development Program (to T.Kas.), Laboratory for Molecular Dynamics of Mental Disorders (to T.Kat.), and Technology and Development Team for Mammalian Genome Dynamics (to K.A.), and Grants-in-Aid from JSPS (KAKENHI 23687010 and 16K14780 to T.Kas.).

## Author Contributions

T.Kas. conceived the study. T.Kas., K.A., and T.Kat. contributed to the design and/or planning of the study. T.Kas. performed all the experiments and bioinformatic analyses, supported by K.A. and T.Kat., except for FISH analysis and search for the *Arse*, which were done by K.M. and A.A, respectively. T.Kas. wrote the manuscript with input from all co- authors.

---

**Figure S1.**
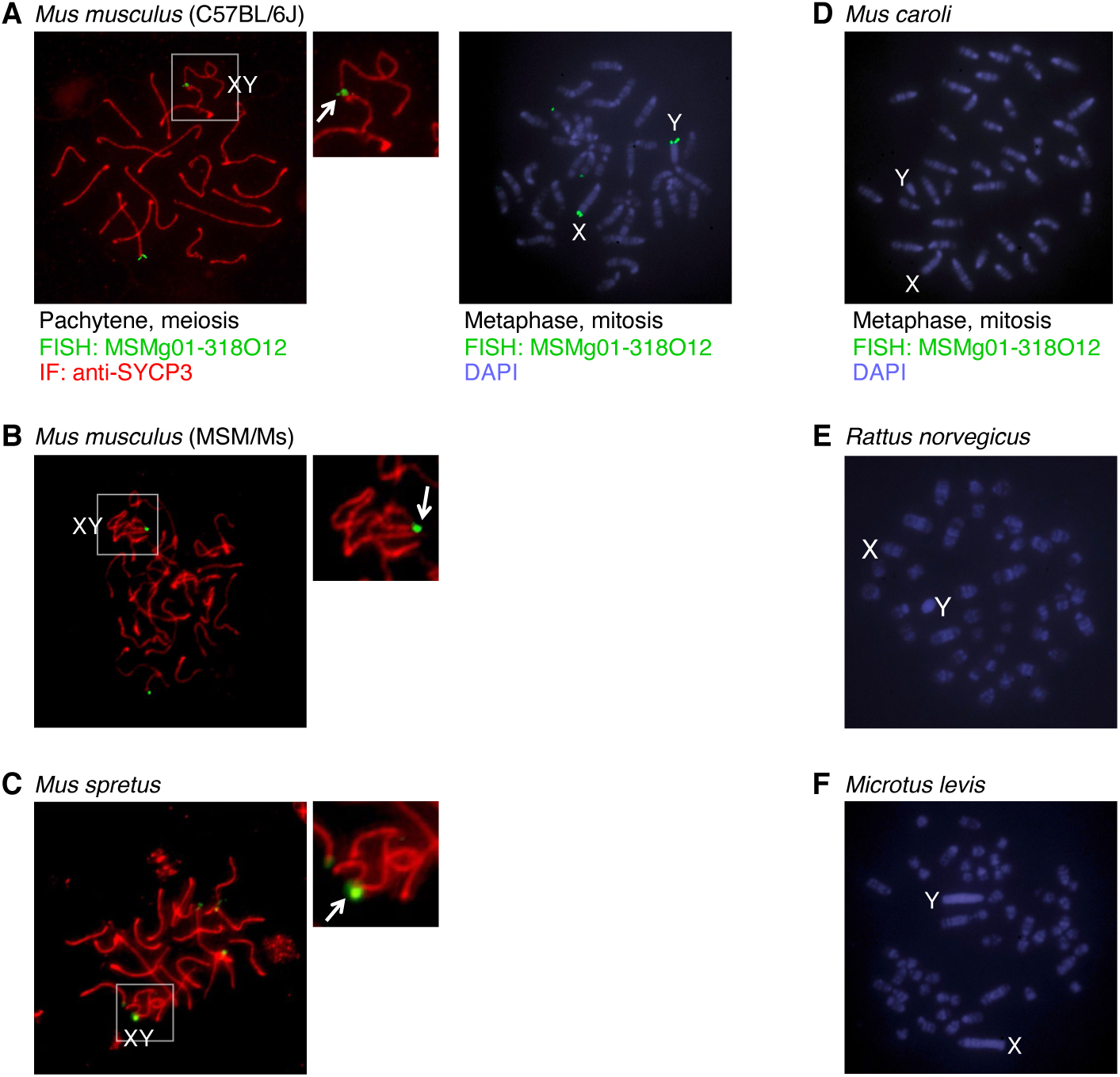
FISH labeling of the PAR- or mo-2-rich regions using the BAC clone (MSMg01-318O12). (**A**) Immunolabeling with FISH (immuno-FISH) for SYCP3 (red) and PAR (green) on pachytene spermatocytes and metaphase FISH analysis of *Mus musculus* (C57BL/6J strain). An arrow indicates the synapsed region of the sex chromosomes (inset). (**B** and **C**) Immuno-FISH analysis of pachytene spermatocytes of *Mus musculus* (MSM/Ms strain) (**B**) and *Mus spretus* (Algerian mouse) (**C**). (**D**–**F**) Metaphase FISH analysis of *Mus caroli* (Ryukyu mouse) (**D**), *Rattus norvegicus* (Rat, F344 strain) (**E**), and *Microtus levis* (East European vole) (**F**).

**Figure S2.**
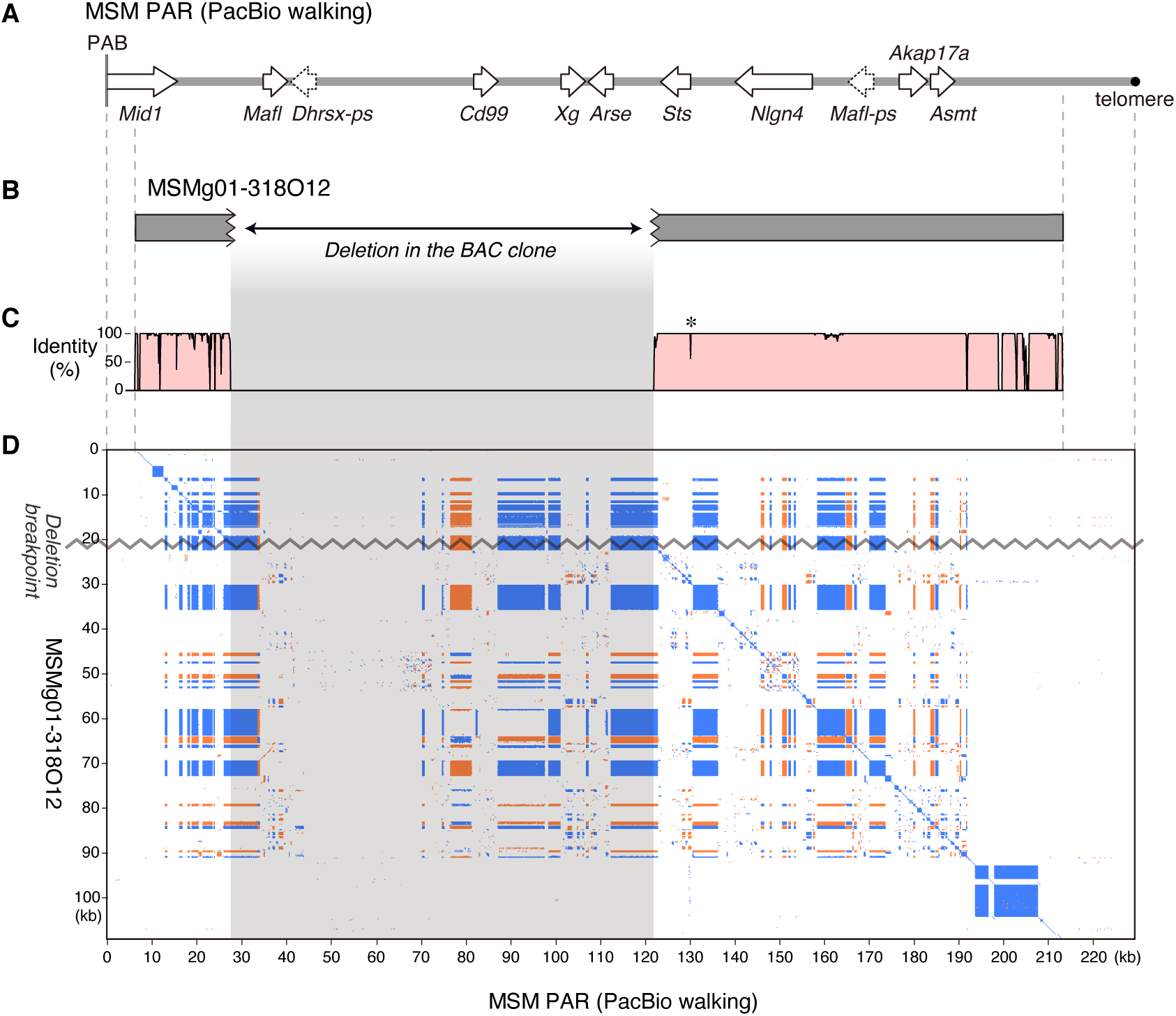
Comparison of the MSM PAR sequence by the PacBio walking method and the BAC clone sequence. (**A**) Gene arrangement in the MSM PAR sequence determined by the PacBio walking method. (**B**) The BAC clone (MSMg01-318O12) with an ∼93-kb deletion. (**C**) DNA sequence identity between the MSM PAR sequence by PacBio walking and the BAC clone calculated using the wgVISTA program (https://genome.lbl.gov/cgi-bin/WGVistaInput). Although most of the differences were found in the repeated sequences, one interesting difference (a 45-bp insertion) was identified in the first exon of the *Sts* gene (an asterisk). This insertion was not found in the MSM BAC clone or the B6J sequence, suggesting the evolution of the MSM PAR during the ∼20 years between the construction of the BAC and the PacBio libraries. (**D**) Dot plots of DNA sequence similarity between the two sequences visualized using the YASS program (https://bioinfo.lifl.fr/yass/yass.php). The deletion in the BAC clone is shown on the horizontal axis (the MSM PAR sequence by the PacBio walking method), and the deletion breakpoint is shown on the vertical axis (the BAC clone sequence).

**Figure S3.**
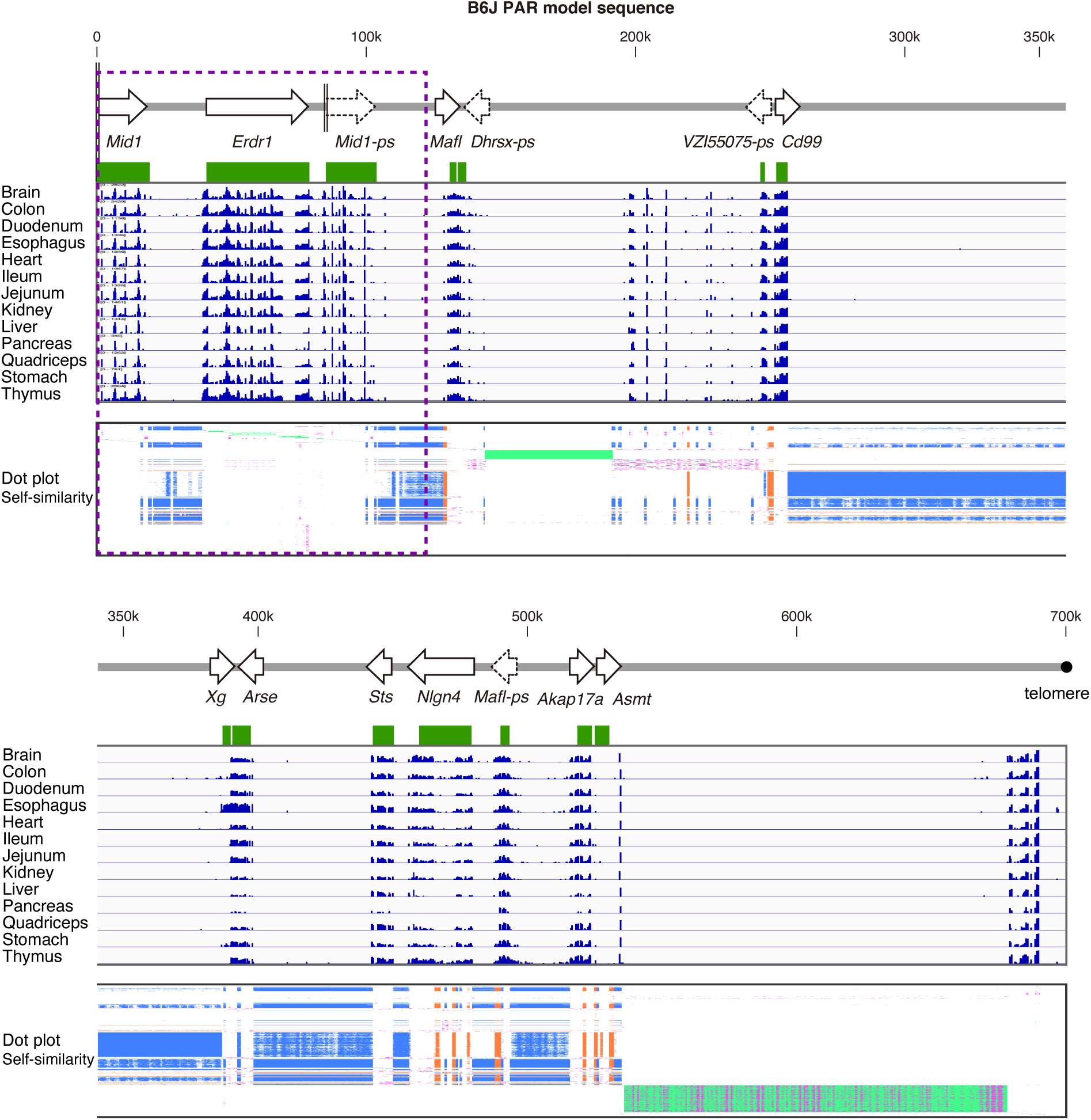
RNA-seq atlas in the mouse PAR. RNA-Seq data from 13 normal tissues of male B6J mice (PRJEB22693) were mapped to the B6J PAR model sequence, and the RNA-seq read density (log_10_) was plotted. The gene arrangement is shown at the top, and the exact locations of the PAR genes/pseudogenes (start codon to stop codon) are indicated by the green rectangles on the RNA-seq read density plots. At the bottom, the dot plot displaying the self-similarity of the B6J PAR sequence (same as Figure 5) with a vertical reduction is shown. The SD region is indicated by a dashed rectangle).

**Figure S4.**
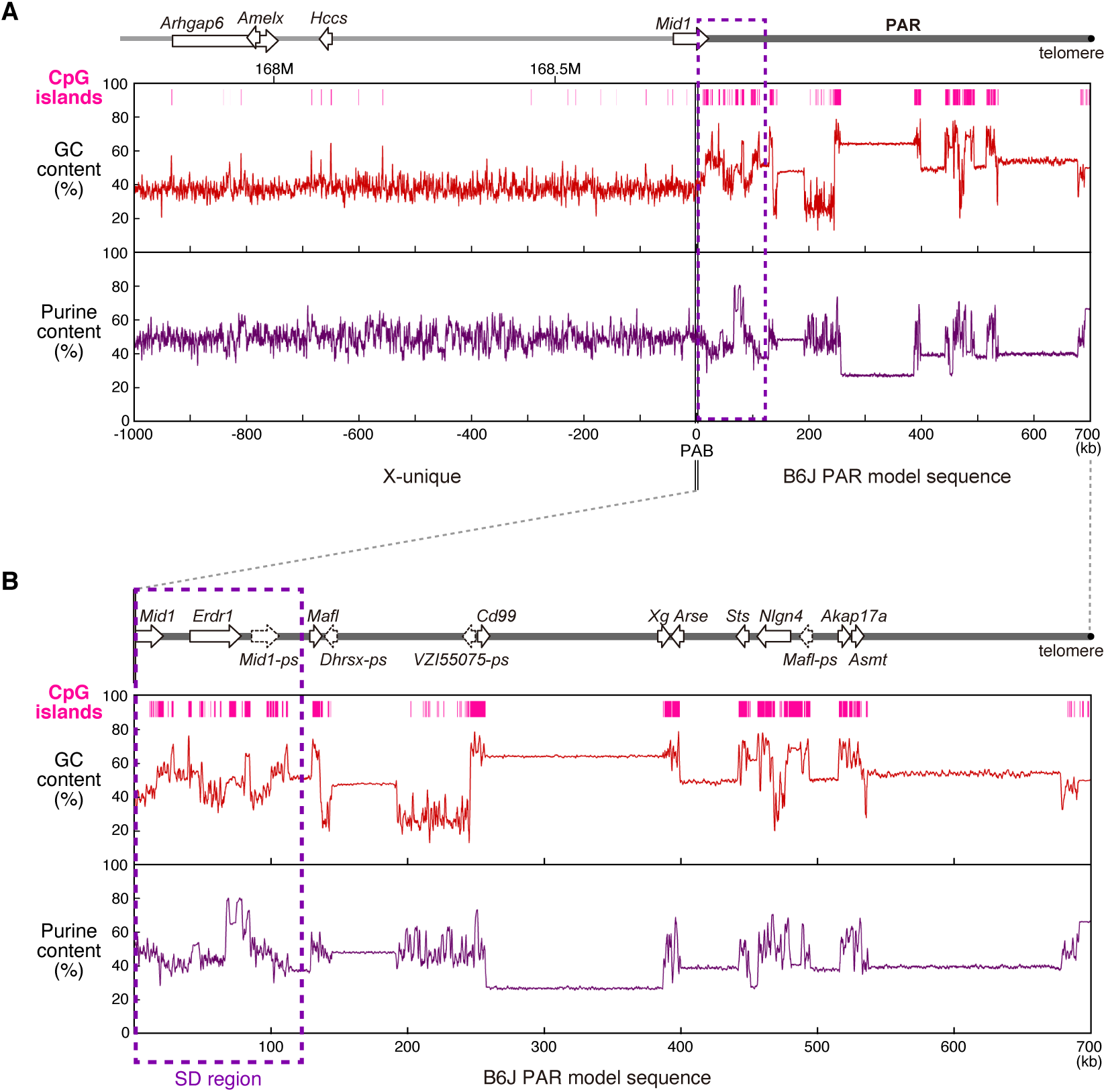
GC and purine contents along the PAB and PAR regions of the mouse X chromosomes. GC (G+C) and purine (A+G) content fluctuated along the 1-Mb PAB region and the PAR (**A**) and in the PAR alone (**B**) measured in 200-bp windows. The gene arrangements and nucleotide positions on the X chromosome (GRCm39) are shown at the top. CpG islands are indicated on the GC content graphs. The SD region is indicated by a dashed rectangle.

**Figure S5.**
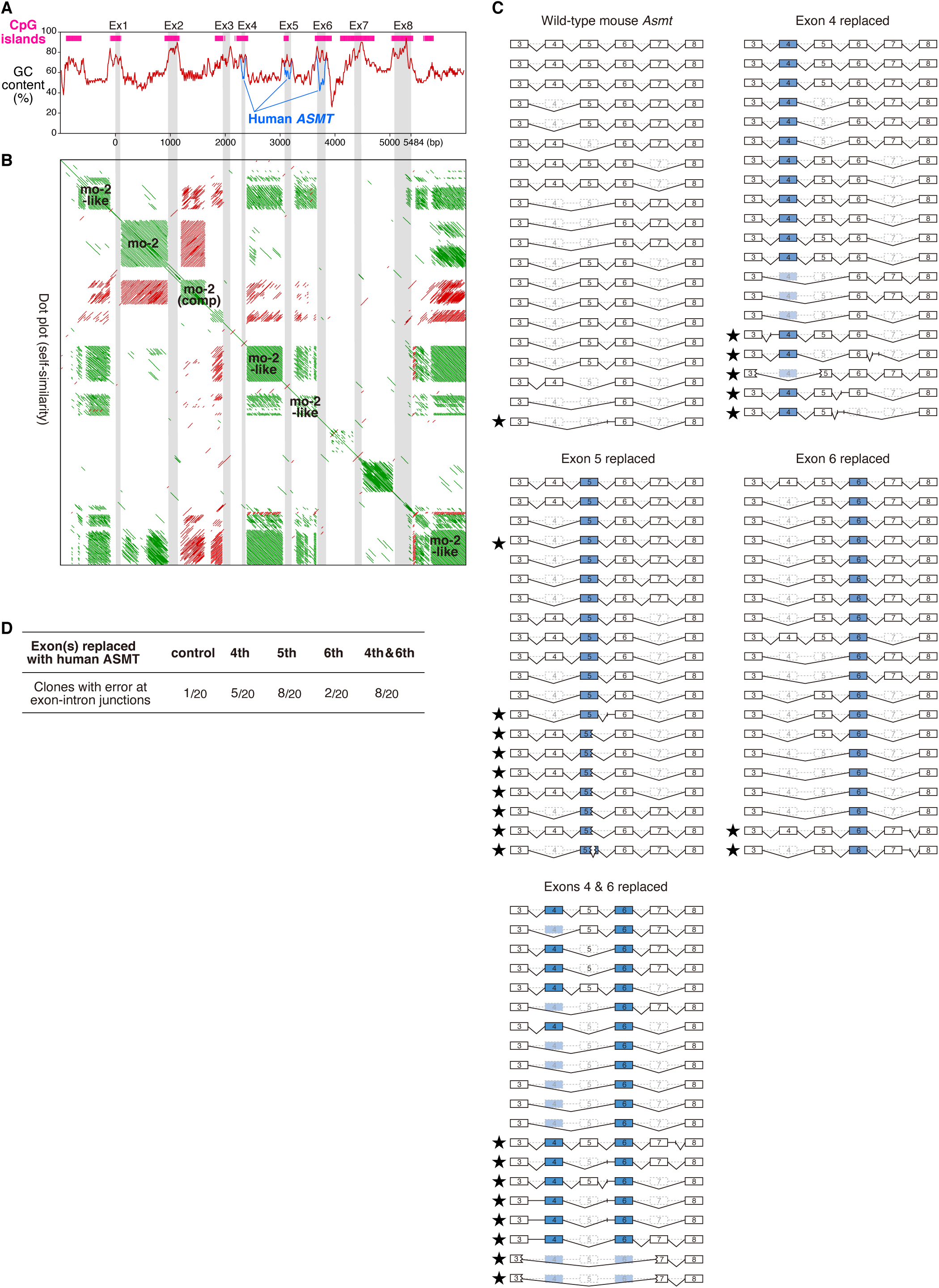
Genomic characteristics of the mouse *Asmt* gene and minigene splicing assay. (**A**) The GC content and predicted CpG islands in the mouse *Asmt* gene region (1,000 bp upstream of the start codon to 1,000 bp downstream of the stop codon). The exons of *Asmt* (Ex 1–8), not including the 5’- or 3’-UTRs, are indicated by gray stripes. The GC content (window size, 50 bp; step size 10 bp) of human *ASMT* exons 4–6, which were used for the minigene splicing assay, is overlaid. (**B**) A dot plot of the self-similarity of the mouse *Asmt* gene. Minisatellites of mo-2, mo-2-like, and complementary mo-2 sequences are shown on the diagonal. (**C** and **D**) Minigene splicing assay. The mouse *Asmt* gene was inserted into a tetracycline-inducible expression vector, and the exon(s) 4, 5, 6, or 4 & 6 were replaced with the corresponding exon(s) of human *ASMT*, which had lower GC content than the mouse exons These minigene constructs were integrated into HEK293FT cells, and stable cell lines were selected. The expression of the minigene was induced, and RNA was extracted. Analysis of transcript splicing was performed on 20 independent *Asmt* mRNA clones. The structures of all the transcripts sequenced are shown (**C**). Abnormal splice donor and acceptor sites were frequently generated (as indicated by stars) in cells carrying the exon-replaced minigene constructs (*P* < 0.05, Fisher’s exact test) (**D**).

**Figure S6.**
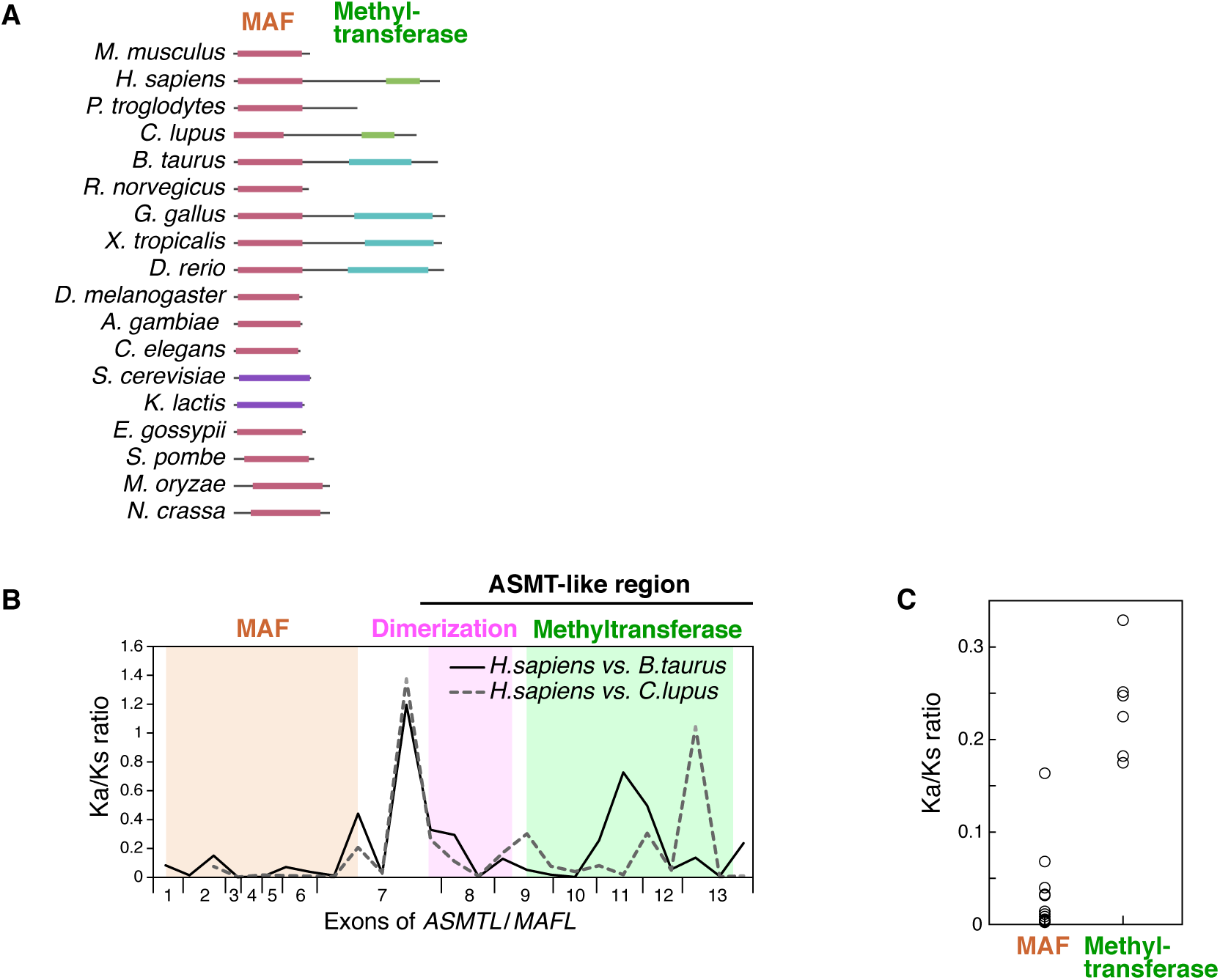
The MAF domain is essential for ASMTL proteins. (**A**) Protein structures of ASMTL homologs among eukaryotes. The schematic drawing was retrieved from HomoloGene (https://www.ncbi.nlm.nih.gov/homologene/?term=asmtl). ASMTL protein, in which the MAF and methylation domains are fused, arose in an autosome prior to the generation of the eutherian PAR, and the ASMTL/MAFL proteins in mouse, rat, and chimpanzee lost the methylation domain. *H. sapiens*, NP_004183.2; *P. troglodytes*, XP_001137782.1; *C. lupus*, XP_005641120.1; *B. taurus*, NP_001030230.1; *R. norvegicus*, NP_001099385.2; *G. gallus*, XP_004938504.1; *X. tropicalis*, NP_001016011.1; *D. rerio*, NP_998676.1; *D. melanogaster*, NP_001027233.1; *A. gambiae*, XP_314812.2; *C. elegans*, NP_492205.2; *S. cerevisiae*, NP_014754.3; *K. lactis*, XP_456133.1; *E. gossypii*, NP_985325.1; *S. pombe*, NP_594969.2; *M. oryzae*, XP_003714242.1; *N. crassa*, XP_964989.1. (**B**) A sliding window analysis of Ka/Ks between human and bovine ASMTL or human and canine ASMTL proteins (window size, 300 bp). Strong purifying selection is observed in the MAF domain. (**C**) Ka/Ks values of the MAF and methyltransferase domains were calculated for all two-pair combinations of ASMTL/MAFL proteins of nine vertebrate species shown in panel **A**. The Ka/Ks ratio was significantly lower in the MAF domain (*P* < 0.0001, *t-*test).

## Supplementary Note

### The mouse PAR gene catalogue

#### Mid1, Mid1-ps

The mouse *Mid1* gene on the X chromosome (NCBI Gene ID: 17318) spans the PAB (see Figure S4) and provides information about the genetic environment both inside and outside the PAR [S1–S3]. Exons 1–3 of the *Mid1* are in the X-unique PAB region, while the rest of the sequence (exon 4 to the last 10th exon) is located inside the PAR. In the B6J strain, each unit of the SD contains the *Mid1* exons 4–10. The same sequence(s) on the Y chromosome is not expressed [S4], so the portions of the *Mid1* gene (exons 4–10) in the SD are likely to be pseudogenes (*Mid1-ps*) except for the *Mid1* that spans the PAB on the X chromosome.

#### Erdr1

No homologs of this gene (Gene ID: 170942) are found in other organisms including other *Mus* species. The gene structure is unusual, in that the entire exon 2 has very high homology with the sequence near the translation start of exon 1, and the amino acid sequences are different due to different reading frames. However, several physiological roles of ERDR1 have been reported [S5–S7]. *Erdr1* deviates from the typical features common to PAR genes that are described later, *i.e*., it has long introns and low GC content at the third position of codons (GC3) (Table 1). Considering that *Erdr1* is not present in the MSM PAR, *Erdr1* must have recently located to the B6J PAR from an autosome. It is possible that the movement of *Erdr1* to the PAR triggered the generation of the SD since the 3’ end of *Erdr1* is just at the border of the SD region.

#### Mafl/Asmtl, Mafl-ps

In humans, the *ASMT* gene and its paralogs (the *ASMTL* genes) are located in PAR1 (Figure 4). ASMTL is a fusion protein composed of almost the entire ASMT and a sequence similar to MAF proteins (Figure S6A). However, the homologs in chimpanzees, rats, insects, plants, and yeast contain only the MAF domain. MAF, named after multicopy- associated filamentation, is a putative inhibitor of septum formation in eukaryotes, bacteria, and archaea [S8]. The human *ASMTL* is expressed in many tissues, but its function is unknown. Examination of the Ka/Ks ratio of *ASMTL* suggested that the MAF domain is more highly negatively selected and is functionally more important than the ASMT domain (Figure 6B,C). In the mouse PAR, there are two sequences (genes) that have homology to MAF. The two have very similar sequences overall; both have no homologous region to ASMT. The former is likely to be a pseudogene for the following reasons: in B6J, the sequence corresponding to the start codon is ATT, and the first ATG is not in the appropriate reading frame; in MSM mice, a premature stop codon appears 24 amino acids after the start of translation. Therefore, we have named the former as *Mafl-ps* and the latter as *Mafl*. Both are expressed in the brain, heart, liver, and kidney of B6J and MSM by RT- PCR.

#### Dhrsx-ps

The human PAR1 contains the *DHRSX* gene, which is widely expressed. Mice have two *DHRSX* homologs, one on chromosome 4 (Gene ID: 236082) and the other on the PAR. As the latter is not expressed based on the mouse EST data and RNA seq data (Figure S3), it is probably a pseudogene (*Dhrsx-ps*).

#### VZI55075-ps

This is a long open-reading-frame (ORF) sequence (corresponding 275 amino acids in B6J) found in *Mus* species (*M. musculus* and *M. caroli* NW_018388668.1). The identical sequence (VZI55075.1; SPRV2_LOCUS23037) was reported in the genome analysis of *Sparganum proliferu* [S9]; however, the genomic DNA of mice, which were the experimental host of the cryptic parasite, probably contaminated the parasite DNA sample. RNA-seq data showed the expression of the ORF sequence (Figure S3), but most EST clones showing significant homology are antisense sequences. Thus, although this ORF sequence has the typical features common to PAR genes (Table 1), this is likely to be a pseudogene in B6J. In MSM, there are many stop codons and frameshifts in the sequence, suggesting that it is already a highly degenerate gene remnant.

#### Cd99

This is one of the two genes that define the XG blood group system, and the human ortholog gene *CD99* is also located inside the PAR1 (Figure 4). A previous paper [S10] reported the cloning of the mouse *Cd99* (NM_025584.2), which was identical to the present B6J PAR sequence, and its location at chromosome 4 shown by FISH analysis. But the gene (Gene ID: 673094) does not map to the autosome in the latest mouse reference sequence, and our experiment measuring the genetic size of the mouse PAR based on recombination frequency (Figure 6A,B) clearly showed that the *Cd99* gene is located inside the PAR.

#### Xg

This is another gene that defines the XG blood group system. The human *XG* gene spans the boundary of PAR1 (Figure 4). Unlike *CD99*, which is widely expressed, the XG protein is present mainly in erythrocytes. RNA-seq analysis showed the expression of mouse *Xg* in the esophagus (Figure S3), though data for erythrocyte was not available. As *Cd99* and *Xg* are located in the mouse PAR, both of those gene products have a number of differences in amino acid sequence between B6J and MSM, which may indicate the presence of a blood group system in mice.

#### Arse

The human protein with the highest sequence similarity to this gene product is arylsulfatase E (HGNC approved gene symbol: *ARSL*), which forms a gene cluster with its paralogs, *ARSD*, *ARSF*, and *ARSH*, in the human X-unique PAB region (Figure 4). Although the gene duplication event generating the arylsulfatase paralogs predated the appearance of the common ancestor of mice and humans [S11], there are no such additional paralogs (*Arsd*, *Arsf*, or *Arsh*) in the mouse genome.

#### Sts

The *Sts* (Gene ID: 20905) was the first mouse PAR gene to be identified [S12]. STS enzyme, steroid sulfatase, is a type of arylsulfatase, and the human *STS* gene is located in the X-unique PAB region, separate from the *ARSD*–*ARSH* gene cluster. In mice, *Sts* and *Arse* exist in the PAR and are next to each other, although there is over 40 kb of repetitive sequences of the mo-2 minisatellite in between (Figure 5).

#### Nlgn4

In humans, the paralogs *NLGN4X* and *NLGN4Y* are located in the X- and Y-unique PAB regions, respectively, whereas in mice, *Nlgn4* is in the PAR without gene duplication. An excellent paper on the mouse *Nlgn4* (Gene ID: 100113365) has recently been published [S13], which reports the rapid evolution of *Nlgn4* and a selective pressure on the protein through a comprehensive sequence analysis of the rodents *Nlgn4* gene. Importantly, *Nlgn4* knockout mice have been generated by gene trapping in embryonic stem cells [S14]. This indicates that may be possible to generate mice deficient in other PAR genes.

#### Akap17a

In human, horse, and dog PARs, the *AKAP17A* gene is next to *ASMT* [S11]. In rats, both *Akap17a* and *Asmt* are located on an autosome but remain located next to each other (NC_051347.1).

#### Asmt

The *Asmt* gene (Gene ID: 107626), which encodes melatonin synthase, is located furthest from the PAB in the mouse PAR (Figure 2). Mutations in the *Asmt* gene have been identified in many mouse strains [S15]. For example, in B6J, two amino acid substitutions (R78G and R242C) result in marked protein instability and loss of the enzymatic activity.

This deprives B6J mice of the pineal hormone melatonin and, interestingly, confers the evolutionary advantage of accelerated gonadal development in a breeding environment [S15, S16].

## References

Abe, K., Nogchi, H., Tagawa, K., Yuzuriha, M., Toyoda, A., Kojima, T., Ezawa, K., Saitou, N., Hattori, M., Sakaki, Y., et al. (2004). Contribution of Asian mouse subspecies *Mus musculus molossinus* to genomic constitution of strain C57BL/6J, as defined by BAC-end sequence-SNP analysis. Genome Res. 14, 2439–2447.

Acquaviva, L., Boekhout, M., Karasu, M.E., Brick, K., Pratto, F., Li, T., van Overbeek, M., Kauppi, L., Camerini-Otero, R.D., Jasin, M., et al. (2020). Ensuring meiotic DNA break formation in the mouse pseudoautosomal region. Nature 582, 426–431.

Amit, M., Donyo, M., Hollander, D., Goren, A., Kim, E., Gelfman, S., Lev-Maor, G., Burstein, D., Schwartz, S., Postolsky, B., et al. (2012). Differential GC content between exons and introns establishes distinct strategies of splice-site recognition. Cell Reports 1, 543–556.

Borodin, P.M., Sablina, O.V., and Rodionova, M.I. (2004). Pattern of X-Y Chromosome Pairing in Microtine Rodents. Hereditas 123, 17–23.

Borodin, P.M., Basheva, E.A., Torgasheva, A.A., Dashkevich, O.A., Golenishchev, F.N., Kartavtseva, I.V., Mekada, K., and Dumont, B.L. (2012). Multiple independent evolutionary losses of XY pairing at meiosis in the grey voles. Chromosome Res. 20, 259–268.

Burgoyne, P.S. (1982). Genetic homology and crossing over in the X and Y chromosomes of mammals. Hum. Genet 61, 85–90.

Dumont, B.L., Williams, C.L., Ng, B.L., Horncastle, V., Chambers, C.L., McGraw, L.A., Adams, D., Mackay, T.F.C., and Breen, M. (2018). Relationship between sequence homology, genome architecture, and meiotic behavior of the dex chromosomes in north american voles. Genetics 210, 83–97.

Flaquer, A., Rappold, G.A., Wienker, T.F., and Fischer, C. (2008). The human pseudoautosomal regions: a review for genetic epidemiologists. Eur. J. Hum. Genet. 16, 771–779.

Fredga, K. (1988). Aberrant chromosomal sex-determining mechanisms in mammals, with special reference to species with XY females. Phil. Trans. R. Soc. Lond. B 322, 83–95.

de la Fuente, R., Parra, M.T., Viera, A., Calvente, A., Gómez, R., Suja, J.Á., Rufas, J.S., and Page, J. (2007). Meiotic pairing and segregation of achiasmate sex chromosomes in eutherian mammals: the role of SYCP3 protein. PLoS Genet. 3, e198.

Hon, T., Mars, K., Young, G., Tsai, Y.-C., Karalius, J.W., Landolin, J.M., Maurer, N., Kudrna, D., Hardigan, M.A., Steiner, C.C., et al. (2020). Highly accurate long-read HiFi sequencing data for five complex genomes. Sci. Data 7, 399.

Honda, T., Suzuki, H., and Itoh, M. (1977). An unusula sex chromosome constitution found in the Amami spinous country-rat, *Tokudaia osimensis osimensis*. Jpn. J. Genet. 52, 247–249.

Jamain, S., Radyushkin, K., Hammerschmidt, K., Granon, S., Boretius, S., Varoqueaux, F., Ramanantsoa, N., Gallego, J., Ronnenberg, A., Winter, D., et al. (2008). Reduced social interaction and ultrasonic communication in a mouse model of monogenic heritable autism. Proc. Natl. Acad. Sci. USA. 105, 1710–1715.

Jensen-Seaman, M.I. (2004). Comparative recombination rates in the rat, mouse, and human genomes. Genome Res. 14, 528–538.

Joseph, A.M., and Chandley, A.C. (1984). The morphological sequence of XY pairing in the Norway rat *Rattus norvegicus*. Chromosoma 89, 381–386.

Kasahara, T., Abe, K., Mekada, K., Yoshiki, A., and Kato, T. (2010). Genetic variation of melatonin productivity in laboratory mice under domestication. Proc. Natl. Acad. Sci. USA. 107, 6412–6417.

Katsura, Y., Iwase, M., and Satta, Y. (2012). Evolution of genomic structures on mammalian sex chromosomes. Curr. Genomics 13, 115–123.

Kipling, D., Salido, E.C., Shapiro, L.J., and Cooke, H.J. (1996). High frequency de novo alterations in the long–range genomic structure of the mouse pseudoautosomal region. Nat. Genet. 13, 78–82.

Lange, J., Yamada, S., Tischfield, S.E., Pan, J., Kim, S., Zhu, X., Socci, N.D., Jasin, M., and Keeney, S. (2016). The landscape of mouse meiotic double-strand break formation, processing, and repair. Cell 167, 695–708.e16.

Matveevsky, S., Kolomiets, O., Bogdanov, A., Hakhverdyan, M., and Bakloushinskaya, I. (2017). Chromosomal evolution in mole coles *Ellobius* (Cricetidae, Rodentia): bizarre sex chromosomes, variable autosomes and meiosis. Genes 8, 306.

Matveevsky, S., Chassovnikarova, T., Grishaeva, T., Atsaeva, M., Malygin, V., Bakloushinskaya, I., and Kolomiets, O. (2021). Kinase CDK2 in mammalian meiotic prophase I: screening for hetero- and homomorphic sex chromosomes. Int. J. Mol. Sci. 22, 1969.

Maxeiner, S., Benseler, F., Krasteva-Christ, G., Brose, N., and Südhof, T.C. (2020). Evolution of the autism-associated *Neuroligin-4* gene reveals broad erosion of pseudoautosomal regions in rodents. Mol. Biol. Evol. 37, 1243–1258.

Palmer, S., Perry, J., Kipling, D., and Ashworth, A. (1997). A gene spans the pseudoautosomal boundary in mice. Proc. Natl. Acad. Sci. USA. 94, 12030–12035.

Pardo-Manuel de Villena, F., and Sapienza, C. (1996). Genetic mapping of *DXYMov15*- associated sequences in the pseudoautosomal region of the C57BL/6J strain. Mamm. Genome 7, 237–239.

Perry, J., and Ashworth, A. (1999). Evolutionary rate of a gene affected by chromosomal position. Curr. Biol. 9, 987–989.

Perry, J., Palmer, S., Gabriel, A., and Ashworth, A. (2001). A short pseudoautosomal region in laboratory mice. Genome Res. 11, 1826–1832.

Peters, A.H.F.M., Plug, A.W., van Vugt, M.J., and de Boer, P. (1997). A drying-down technique for the spreading of mammalian meiocytes from the male and female germline. Chromosome Res. 5, 66–68.

Raudsepp, T., and Chowdhary, B.P. (2015). The eutherian pseudoautosomal region. Cytogenet. Genome Res. 147, 81–94.

Roy, M., Kim, N., Xing, Y., and Lee, C. (2008). The effect of intron length on exon creation ratios during the evolution of mammalian genomes. RNA 14, 2261–2273.

Salido, E.C., Li, X.M., Yen, P.H., Martin, N., Mohandas, T.K., and Shapiro, L.J. (1996). Cloning and expression of the mouse pseudoautosomal steroid sulphatase gene (*Sts*). Nat. Genet. 13, 83–86.

Sarsani, V.K., Raghupathy, N., Fiddes, I.T., Armstrong, J., Thibaud-Nissen, F., Zinder, O., Bolisetty, M., Howe, K., Hinerfeld, D., Ruan, X., et al. (2019). The genome of C57BL/6J “Eve”, the mother of the laboratory mouse genome reference strain. G3 9, 1795–1805.

Schilthuizen, M., Giesbers, M.C.W.G., and Beukeboom, L.W. (2011). Haldane’s rule in the 21st century. Heredity 107, 95–102.

Söllner, J.F., Leparc, G., Hildebrandt, T., Klein, H., Thomas, L., Stupka, E., and Simon, E. (2017). An RNA-Seq atlas of gene expression in mouse and rat normal tissues. Sci. Data 4, 170185.

Takahashi, Y., Mitani, K., Kuwabara, K., Hayashi, T., Niwa, M., Miyashita, N., Moriwaki, K., and Kominami, R. (1994). Methylation imprinting was observed of mouse mo-2 macrosatellite on the pseudoautosomal region but not on chromosome 9. Chromosoma 103, 450–458.

Zhang, C., Clough, S.J., Adamah-Biassi, E.B., Sveinsson, M.H., Hutchinson, A.J., Miura, I., Furuse, T., Wakana, S., Matsumoto, Y.K., Okanoya, K., et al. (2021). Impact of endogenous melatonin on rhythmic behaviors, reproduction, and survival revealed in melatonin-proficient C57BL/6J congenic mice. J. Pineal. Res. 71, e12748.

## Supplementary References

S1 Palmer, S., Perry, J., Kipling, D. & Ashworth, A. A gene spans the pseudoautosomal boundary in mice. Proc Natl Acad Sci USA 94, 12030–12035 (1997).

S2 Perry, J. & Ashworth, A. Evolutionary rate of a gene affected by chromosomal position. Curr Biol 9, 987–989 (1999).

S3 Montoya-Burgos, J. I., Boursot, P. & Galtier, N. Recombination explains isochores in mammalian genomes. Trends Genet 19, 128–130 (2003).

S4 Lancioni, A., et al. Lack of *Mid1*, the mouse ortholog of the Opitz syndrome gene, causes abnormal development of the anterior cerebellar vermis. J Neurosci 30, 2880–2887 (2010).

S5 Houh, Y., Kim, K., Park, H. & Cho, D. Roles of erythroid differentiation regulator 1 (Erdr1) on inflammatory skin diseases. Int J Mol Sci 17, 2059 (2016).

S6 Weis, A. M., Soto, R. & Round, J. L. Commensal regulation of T cell survival through Erdr1. Gut Microbes 9, 458–464 (2018).

S7 Abo, H., et al. Erythroid differentiation regulator-1 induced by microbiota in early life drives intestinal stem cell proliferation and regeneration. Nat Commun 11, 513 (2020).

S8 Butler, Y. X., Abhayawardhane, Y. & Stewart, G. C. Amplification of the *Bacillus subtilis maf* gene results in arrested septum formation. J Bacteriol 175, 3139–3145 (1993).

S9 Kikuchi, T., et al. Genome of the fatal tapeworm *Sparganum proliferum* uncovers mechanisms for cryptic life cycle and aberrant larval proliferation. Commun Biol 4, 649 (2021).

S10 Park, S. H., et al. Rapid divergency of rodent CD99 orthologs: Implications for the evolution of the pseudoautosomal region. Gene 353, 177–188 (2005).

S11 Katsura, Y., Iwase, M. & Satta, Y. Evolution of genomic structures on mammalian sex chromosomes. Curr Genomics 13, 115–123 (2012).

S12 Salido, E. C., et al. Cloning and expression of the mouse pseudoautosomal steroid sulphatase gene (*Sts*). Nat Genet 13, 83–86 (1996).

S13 Maxeiner, S., Benseler, F., Krasteva-Christ, G., Brose, N. & Südhof, T. C. Evolution of the autism-associated *Neuroligin-4* gene reveals broad erosion of pseudoautosomal regions in rodents. Mol Biol Evol 37, 1243–1258 (2020).

S14 Jamain, S., et al. Reduced social interaction and ultrasonic communication in a mouse model of monogenic heritable autism. Proc Natl Acad Sci USA 105, 1710– 1715 (2008).

S15 Kasahara, T., Abe, K., Mekada, K., Yoshiki, A. & Kato, T. Genetic variation of melatonin productivity in laboratory mice under domestication. Proceedings of the National Academy of Sciences 107, 6412–6417 (2010).

S16 Zhang, C., et al. Impact of endogenous melatonin on rhythmic behaviors, reproduction, and survival revealed in melatonin-proficient C57BL/6J congenic mice. J Pineal Res 71, e12748 (2021).

